# Climatic variations along an aridity gradient drive significant trait intraspecific variability in Mediterranean species

**DOI:** 10.1101/2022.08.21.504686

**Authors:** Lorenzo Maria Iozia, Virginia Crisafulli, Laura Varone

## Abstract

Summer drought represents one of the main stress sources stress for plant communities in the Mediterranean region. Plants can adopt several response strategies to cope with stress, reflected in the adoption of specific Plant Functional Traits (PFTs). Trait-based approaches commonly meet three critical issues: they overlook Intraspecific Variability (ITV), they focus on a large spatial scale, or they focus on single trait responses to stress. In this study, we present evidence for a significant amount of ITV in morphological and anatomical trait syndromes observed between three local populations of *Phyllirea latifolia, Pistacia lentiscus* and *Quercus ilex,* distributed along an aridity gradient. Thicker, more physiologically expensive leaves and lower heights found in the drier sites mainly conform to drought-resistance strategies. Interestingly, PFTs from *Cistus salviifolius* were found not to vary between sites. This implies that not all species vary at the same geographical scale, possibly depending on their different successional role. The main implication behind our findings is that climate can easily drive significant ITV in multiple traits among plant populations, even at a local scale, although trait responsiveness might be species-specific. Different plant populations hailing from the same geographical regions might thus respond differently to climate change.

**HIGHLIGHT:** Variations in Plant Functional Traits from several Mediterranean species found along an aridity gradient on a local (<60Km radius) scale; responses consistent to reported drought adaptations.

## INTRODUCTION

Plant Functional Traits (PFTs) are known to affect plant performance through their effects on functional processes such as growth, survival, and reproduction (Violle *et al*., 2007). PFTs are thus very often employed in the study of plant adaptive strategies, which according to trait–based ecology ultimately results from specific trait combinations (Zhou *et al*., 2022). In this view, trait covariation described by the Leaf Economic Spectrum (Wright *et al*., 2004) identified a global pattern of leaf trait variation along wide environmental gradients, contributing to clarify the mechanism underlying plant responses to environmental changes (Maharjan *et al*., 2021). Over the last decade this approach was further developed by Reich (2014), who also included water as a key resource to the framework and integrated stems and roots to the spectrum of traits to help explain plant conservative and acquisitive resource uptake strategies.

An important issue of studies embracing trait-based approaches is that they mainly focus on interspecific, or between-species, variability (BTV) (Albert *et al*., 2011; Funk *et al*., 2017). This can be relevant as, albeit often being a lesser driver of variability, several evidence suggests a significant effect of intraspecific variability (ITV) on several ecological processes (Albert *et al*., 2011; Puglielli *et al*., 2022) such as disease resistance (Garrett *et al*., 2009), decomposition rates (Lecerf and Chauvet, 2008), community assemblies (Cornwell *et al*., 2008; Jung *et al*., 2010) and response to precipitation and temperature (Jung *et al*., 2014; Solé-Medina *et al*., 2022; Rodríguez-Alarcón *et al*., 2022).

Spatial scale also becomes a relevant issue when tackling ITV, as the most intuitive approach to observe variations among species would be to confront populations very distant from each other. This approach logically ensures to cover a wide range of conditions to maximize the amount of ITV observed. However, spatial environmental heterogeneity is more likely to drive genetic divergence than distance between populations (Hufford and Mazer, 2003).

Another issue arises in studies that focus on single or few traits’ responses to drought: multiple response strategies might involve several traits simultaneously, leading to possibly confusing results (Rodríguez-Alarcón *et al*., 2022). Therefore, it is important to consider responses within a multivariate functional space, even better if covering multiple plant organs, to understand how coordinated trait shifts might lead to alternative strategies against drought (Blumenthal *et al*., 2020).

The aim of this study is to answer the question: *could climate variations significantly affect plant functional traits at a local scale?* Recent evidence suggests that climate-driven ITV can indeed be observed at a local scale (Moreira *et al*., 2012; Kumordzi *et al*., 2014; Garcia *et al*., 2022; Dalla Vecchia and Bolpagni, 2022), but there is still a substantial scarcity of studies regarding this issue. This paucity is accentuated by the ambiguous definition of locality throughout literature: *local* can arbitrarily point to different spatial scales, from the shores of a small, shallow lake (Dalla Vecchia and Bolpagni, 2022) to small archipelagos scattered over hundreds of Kilometres (Moreira *et al*., 2012; Kumordzi *et al*., 2014). Given that the most used high resolution climate models are made available at 30 arc seconds (Fick and Hijmans, 2017; Karger *et al*., 2017), for the purpose of this study we considered populations *local* when situated in a ~60 Km radius from a given point.

To investigate local scale intraspecific variations, we studied the variation of trait syndromes including several plant functional traits from a selection of plants often found cohabiting in the Mediterranean shrubland. To inspect the effects of climate as a natural selector, we focused on populations chosen along an aridity gradient, as aridity is among the most important selective factors driving the evolution of plants’ adaptive strategies in the Mediterranean region (Basu *et al*., 2016). At the same time, the Mediterranean region shows a very high environmental heterogeneity, offering a wide range of climatic variations at a local scale (Lionello *et al*., 2006). Consistently with trait-based economic spectrum models, populations of Mediterranean species tend to adopt conservative acquisition strategies when inhabiting more xeric habitats, while populations in more mesic environments are rewarded by acquisitive strategies (Solé-Medina *et al*., 2022). The dichotomy between conservative and acquisitive resource uptake strategies is also reflected in BTV, with conservative species appearing more drought resistant and showing more ITV than acquisitive species (Rodríguez-Alarcón *et al*., 2022).

To answer our question, we compared a combination of plant functional traits known to be highly responsive to aridity. Indeed, plants typically adopt thicker, smaller leaves when adapted to dry environments. This trend is reflected in the way different functional traits respond to drought: Leaf Dry Matter Content (LDMC) shows consistent increases with increasing aridity (Anderegg *et al*., 2021), much like Shoot Length Growth Efficiency (LE) (Crescente *et al*., 2002). Leaf Mass per Area (LMA) usually increases along aridity gradients (Wright *et al*., 2005; Ivanov *et al*., 2008; Anderegg *et al*., 2021), although annual plants exhibiting a drought escape strategy have been reported adopting lower LMA values in drier conditions (Welles and Funk, 2021). Similar contrasting responses can be seen for Stomatal Density (SD) as well, either with increases (Carlson *et al*., 2016) or decreases (Guo *et al*., 2017) in response to aridity. Furthermore, some traits can exhibit inverse correlation with precipitation, such as Mean Plant Height (H) (Nunes *et al*., 2017) and wood density (i.e., Specific Stem Density, SSD) (Martínez-Cabrera *et al*., 2009). Leaf anatomy can also be affected at a microscopic scale, as plants living in arid conditions also show thicker epidermis and denser tissue, often with abundant sclerenchyma and reduced spongy mesenchyme (Dörken *et al*., 2020).

Given these premises, the main hypothesis behind this study is that plant populations living in close proximity but exposed to different climate conditions will adopt different functional trait syndromes to embrace appropriate response strategies to their local environment. More precisely, we expect plants from drier sites to significantly differ from their counterparts hailing from wet sites, adopting conservative strategies reflected in higher LDMC, LE and LMA; lower SD, H, SSD and thicker leaf tissues. In agreement to literature, we also expect conservative species to present greater ITV in response to drought than species known to generally adopt more acquisitive strategies (Rodríguez-Alarcón *et al*., 2022).

## MATERIALS AND METHODS

### Study sites

The study was carried out in central Italy, spanning over an area of approximately 1600 Km^2^ in the *Latium* region, (Fig. 1) during the period June-September 2021. Climate strongly varies both spatially and temporally in this region, allowing for the identification of several phytoclimatic regions, from temperate to Mediterranean, commonly subjected to various degrees of summer drought (Blasi, 1994).

**Figure 1.**
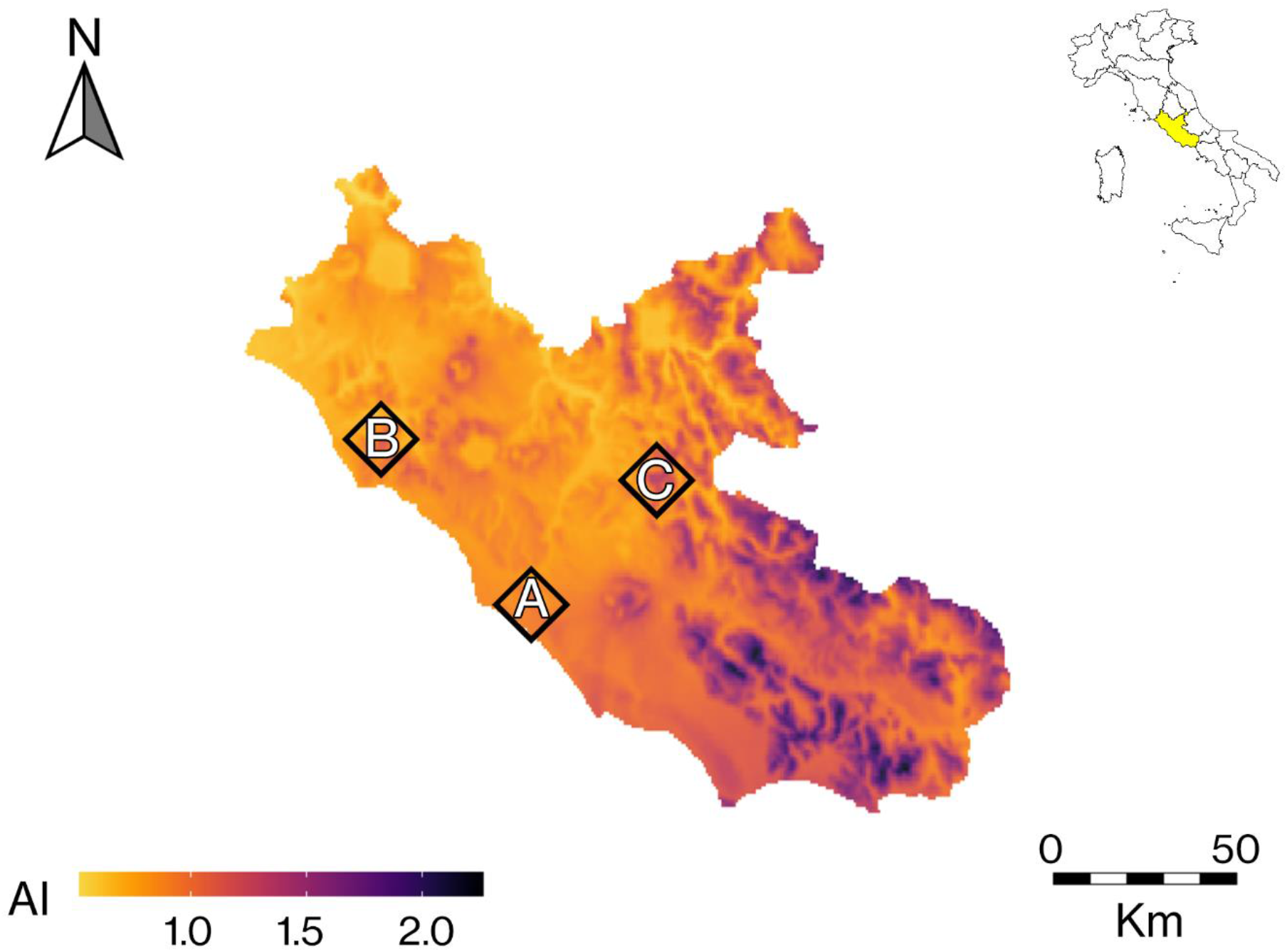
Aridity Index Map of the *Latium* region. Study sites are labelled on the map as **A** for Castel Fusano (RM), **B** for La Farnesiana (RM) and **C** for Monte Catillo in Tivoli (RM). An estimation of the Aridity Index is presented by the colour gradient, from yellow (AI=0) to blue (AI=2.3). The *Latium* region is highlighted by the colour yellow on the map of Italy.

Three sampling sites of 1 Km^2^ were chosen along an aridity gradient with a mean distance from each other of 64.6 Km: site A located at *Castel Fusano* (3 m asl, 41°43’23.6 “N, 12°19’55.7” E), site B at *Tenuta La Farnesiana* (150 m asl, 42°11’38.9 “N, 11°52’33.1” E), and site C at the *Monte Catillo* Natural Reserve (430 m asl, 41°57’51.5 “N, 12°48’54.9” E). Over the last decade, Site A (Fig. 2A) was characterised by a mean temperature of 16.0 °C; a mean maximum temperature of 22.3 °C, and a mean minimum temperature of 9.9 °C. Total mean rainfall in site A was 847.7 mm. This site was found to be the driest of the three chosen sites. At site B (Fig. 2B) the mean temperature was 15.1 °C, the mean maximum temperature 21.4 °C, and the mean minimum temperature 9.5 °C. Mean total annual rainfall was 938.0 mm. The site was therefore characterised by an intermediate aridity level. Site C (Fig. 2C) was characterised by a mean annual temperature of 13.9 °C, a mean maximum temperature of 22.1 °C, and a mean minimum temperature of 8.1 °C. The mean total rainfall was 1306.1 mm. This site was thus defined as the wet site.

**Figure 2.**
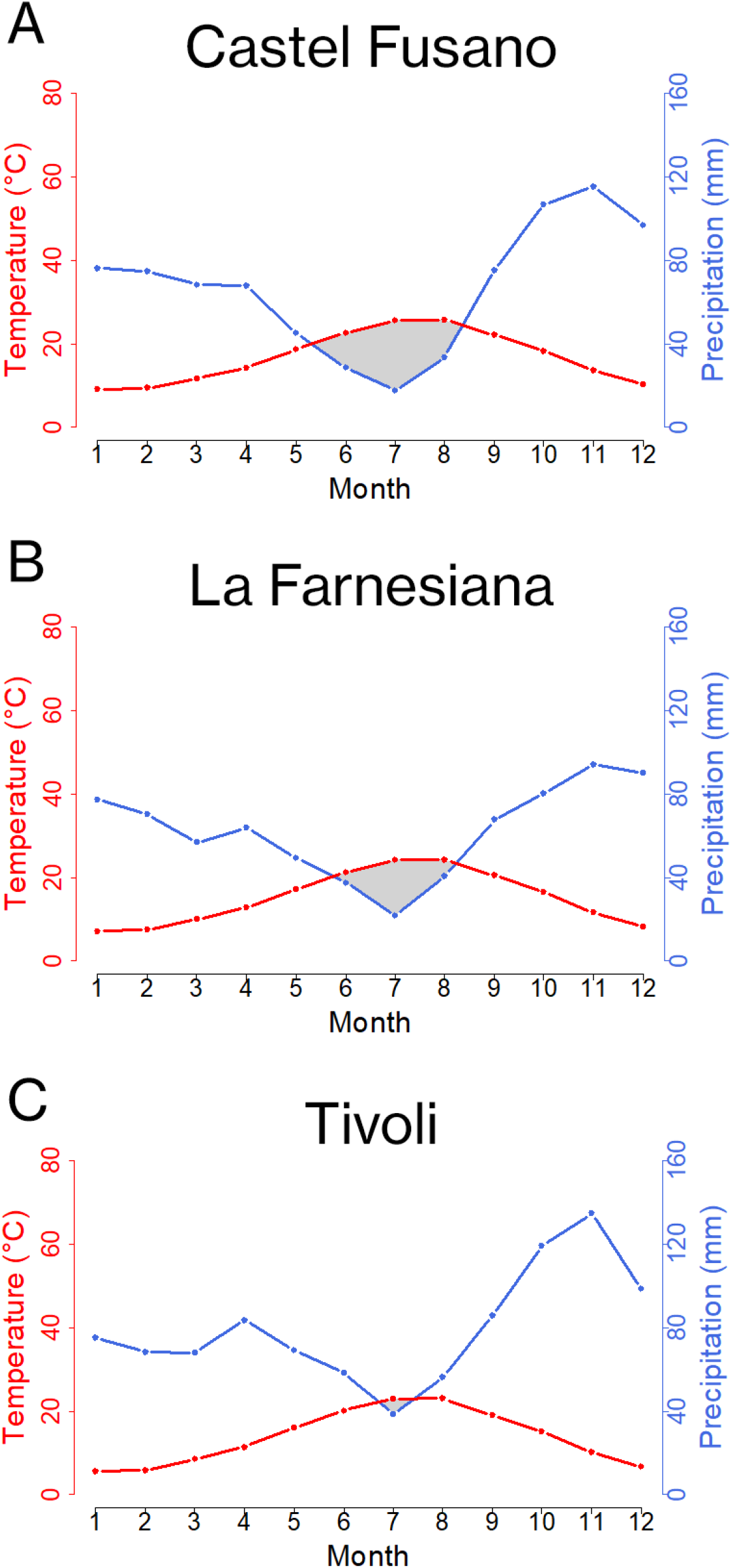
Ombrothermic diagrams of our study sites obtained from the ChelsaClimate model (Karger *et al*., 2017), according to Bagnouls and Gaussen (1953). Site **A** is Castel Fusano (RM), site **B** La Farnesiana (RM) ad site **C** Monte Catillo in Tivoli (RM).

Local climatic data were explored using the ChelsaClimate model (Karger *et al*., 2017) and then confirmed with data provided by the nearest Meteorological Stations (‘SIARL-Servizio Integrato Agrometeorologico della Regione Lazio’, 2022): *Fiumicino - Maccarese* (3 m asl, 41° 49’11.2 “N, 12° 14’47.5” E) for site A, *Blera - Puntoni* (258 m asl, 42° 16’07.6 “N, 12° 00’58.1” E) for site B, and *Licenza - Colle Franco* (460 m asl, 42° 03’56.7 “N, 12° 54’39.9” E) for site C; all referring to the period 2011-2021. Aridity was quantified through the combined use of the FAO-UNEP Aridity Index (UNEP, 1992) and Umbro thermal Diagrams generated according to Bagnouls and Gaussen (1953).

Moreover, soil samples were collected at each site in September 2021, with a manual drill, at 40 cm depth. Five 1 kg soil samples were collected at each sampling site. Analyses were performed by a specialised laboratory, according to the official methodologies of the National Pedological Observatory of Agricultural, Food and Forest Resources Ministry (Jones Jr, 1999).Data on soil characteristics are showed in Appendix S1.

### Study species

The study was carried out on 4 co-occurring Mediterranean plant species, chosen for their simultaneous presence in each sampling site and diverse functional type and evolutive origin: *Quercus ilex* L., *Cistus salviifolius* L., *Phyllirea latifolia* L., and *Pistacia lentiscus* L. The former two are widely considered indigenous taxa evolved in relatively recent time under the Mediterranean climate (Correia and Catarino, 1994; Blondel and Aronson, 1999), although some debate has recently sparked over Holm oak’s origin (Martín-Sánchez *et al*., 2022). The other species represent pre-Mediterranean elements: *Phillyrea* is an Afro-tropical taxa originated from arid sites (Quézel, 1985), and *Pistacia* evolved in the semi-arid steppes of Central Asia (Blondel and Aronson, 1999). Pre-Mediterranean and Mediterranean taxa are known to attain drought resistance by different traits or combination of traits; nevertheless, the extent of these adaptations also varies considerably among species co-occurring in the same environments (Gratani and Bombelli, 2001; Pesoli *et al*., 2003; Gratani and Varone, 2004).

In addition, the selected species also included two main functional types: evergreen sclerophyllous (Q. *ilex, P. lentiscus* and *P. latifolia)* and drought semi-deciduous (C. *salvifolius),* which adopt diverse adaptive strategies (Harley *et al*., 1987; Stefi and Christodoulakis, 2021; Martín-Sánchez *et al*., 2022; Vignola *et al*., 2022) strongly differing in terms of structural and physiological leaf traits and drought resistance (Gratani *et al*., 2018). More specifically, evergreen sclerophylls tend to adopt conservative strategies such as thicker leaves and deeper roots, while *C. salviifolius* tends to avoid drought by shedding a high percentage of leaves every four to eight months, also adopting thicker leaves during summer, adapting to drier conditions, and acting as a pioneer species in succession towards woodland (Gratani and Crescente, 1997; Grant *et al*., 2014).

### Experimental design

Each sampling site was divided in five plots of 100 m^2^, each containing at least four individuals for each species. For each plot and for each species, four healthy adult individuals were chosen for measurements. Between all species, this adds up to 240 individuals. We randomly collected three fully expanded sun leaves and one shoot from each specimen, harvesting them from the upper part of the crown in the southern side of the plant to minimize variations in sun exposure. Leaves were stored in dark, refrigerated, and moist conditions according to Pérez-Harguindeguy *et al.* (2016) and transported to Sapienza University of Rome to carry out our analyses. Shoots were collected at the end of the growing season in July, to measure their maximum length (cm), and stored with the same procedure as the leaves.

### Leaf and Whole Plant Functional traits

For each species, we measured the plant height (H, cm) of the selected individuals by using a digital clinometer (Haglöf, Sweden). For each site and for each species, leaf morphological analysis was conducted on randomly chosen, fully expanded leaf samples (n = 20). The following parameters were measured for each leaf: fresh leaf area (LA, cm^2^), measured using the image analysis system Delta-T Devices (UK); leaf water saturated mass (SM, mg), measured after rehydration until saturation for 48 h at 5°C in the darkness; leaf dry mass (DM, mg), obtained by drying the leaves in a thermostatic oven (M710 Thermostatic Oven) at 90 °C until constant weight was reached.

We then used our measurements to produce the following parameters: leaf mass per area (LMA, mgcm^-2^), given by the ratio between DM and LA (Larcher, 2003); leaf dry matter content (LDMC, mgg^-1^), given by the ratio between DM and SM (Pérez-Harguindeguy *et al*., 2016).

### Stem and Shoot Functional traits

A total of 240 representative shoot samples, one for each selected specimen, were separated into leaf and stem mass to measure leaf surface area (LA_S_, cm^2^), leaf dry mass (DM_S_, mg), and stem dry mass (SDM_s_, mg), according to the same methods mentioned above. Each stem was measured with a meter to obtain stem length (SL_s_, mm), while stem diameter (SD_s_, mm) was measured using a Digital Micrometre (Kennedy 331-301, 0-1 ″, 0-25 mm). Fresh stem volume (SV_s_, mm^3^) was estimated approximating the stem to a cylinder, using the formula:

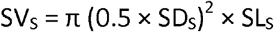

Measurements were then used to produce the following parameters: specific stem density (SSD, gem^-3^) given by the ratio between SDM_s_ and SV_s_ (Pérez-Harguindeguy *et al*., 2016); shoot length growth efficiency (LE, cm^3^g^-1^), calculated according to King (1997) and Crescente *et al.* (2002) as:

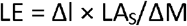

where Δl was the shoot length growth increment, calculated at the end of the growing season, LAs the total leaf area per shoot, and ΔM the total dry mass (leaf + stem) per shoot.

### Anatomical Leaf Functional traits

A total of 120 representative fresh leaf samples (n=10 for each site for each species) were selected to carry out anatomical analyses. Each leaf was dissected to half its length and the sections were dehydrated in a 90% ethanol aqueous solution. Sections were observed with a Nikon Eclipse E400 optical microscope (Hall *et al*., 2013) and digitally analysed (AxioVision AC, Release 4.5). All measurements were restricted to free-vein lamina areas (Chabot and Chabot, 1977). The following parameters were measured: thickness of the abaxial leaf epidermis (AE, μm); palisade parenchyma thickness (_PAL_PT, μm); and patchy parenchyma thickness (_PAT_PT, μm).

From these parameters was calculated the ratio between palisade and patchy parenchyma thickness (PP), according to Ferreira De Melo Junior and Boeger (2016).

Abaxial Stomatal Density (SD, n° of stomata*mm^-2^) was also determined using the method of transparent polish impressions on the lower leaf page, according to Sack *et al.* (2003). Gel impressions, obtained from 10 sun leaves for each population coming from different individuals, were observed with an optical microscope to determine the number of stomata in a 220×165 μm^2^ area.

### Statistical analysis

To detect variations among functional traits, we performed a One-way ANOVA to analyse differences between trait means from each site. Tukey’s test was then used to allow for multiple comparisons.

Variance partitioning analysis (VPA) was used to quantify the amount of variation explained by each variable (Šmilauer and Lepš, 2014). In this specific case, we quantified the amount of variability expressed between species and among species basing on their proportional unique contribution to total variability in our dataset. The likelihood of species and site factors in explaining the total variance of functional traits was determined, according to de Bello *et al.* (2011). Significance of each component of the VPA model was also evaluated performing Monte Carlo permutation test of the predictor effect.

To analyse trait syndromes, a Principal Component Analysis (PCA) based on the correlation matrix was performed. The analysis included the following parameters: H, SSD, SD, PP, AE, LE, LMA, LDMC. PCA analysis were performed separately for each species, in order to reduce noise from BTV when comparing different sites. Eigen analysis on varimax-rotated principal components and Pearson pair-wise correlation coefficients between the first two principal components and original trait parameters were then used to characterise bivariate relationships between PCs and traits.

To address our main question, Principal Components were then statistically analysed using Trait Probability Distributions analysis (TPD) and multiple Wilcoxon-Mann-Whitney U tests. The first analysis allows to confront variables by directly assessing the amount of dissimilarity among their respective probability distributions (Carmona *et al*., 2019), while the Wilcoxon-Mann-Whitney U test was chosen for its ability to identify statistically significant differences among non-parametrical distributions. For this last analysis, differences were considered significant for p ≤ 0.05.

Statistical data processing was carried out using the R 4.0.3 statistical analysis software (The R Foundation for Statistical Computing, 2019); VPA was performed using the R package vegan 2.5-7 (Oksanen *et al*., 2013); PCA was performed with the R package ade4 1.7-16 (Thioulouse *et al*., 2018); TPD was performed using the R package TPD 1.1.0 (Carmona *et al*., 2019).

## RESULTS

### Intraspecific variability in plant functional traits

Almost all considered species showed marked differences in their PFTs among study sites, with *C. salviifolius* being the species showing the least amount of ITV with the only significant (p < 0.05) difference across sites in LMA (Table S2). The PFTs that showed the most significant (p < 0.05) variations among all species were H and LMA, with LDMC, SSD, LE and AE varying less frequently. SD was the least variable trait, showing divergences only for *P. latifolia.* Mean (±ES) (n = 20) PFTs values and variations are presented in Appendix 2.

### Data processing and statistical analysis

VPA showed that both species (0.508) and sites (cumulative effects of sites A, B and C) (0.113) had a significant effect on variance (p < 0.05), with a greater explanatory power from species. The combined effect of these factors was not significant, while the unexplained part of total variance was 0.403. Given the larger weight of BTV on total variability, species had consequently been separated and presented individually to better appreciate the effect of ITV.

### *Quercus ilex* L

PCA on Q. *ilex* (Fig. 3A) returned two principal components explaining the greater variation of the traits, amounting to 63% of the total variance (35.8% by PC1 and 27.2% by PC2). PC1 was significantly related (p < 0.05) to AE and SSD and inversely to LDMC, LE, SD and PP, while PC2 was significantly related to H, SD, and PP and inversely to LMA (Fig. 3B). Populations occupied different dials, with site A showing higher values of LMA, LDMC, LE, SD and PP and lower H when compared to the other sites.

**Figure 3.**
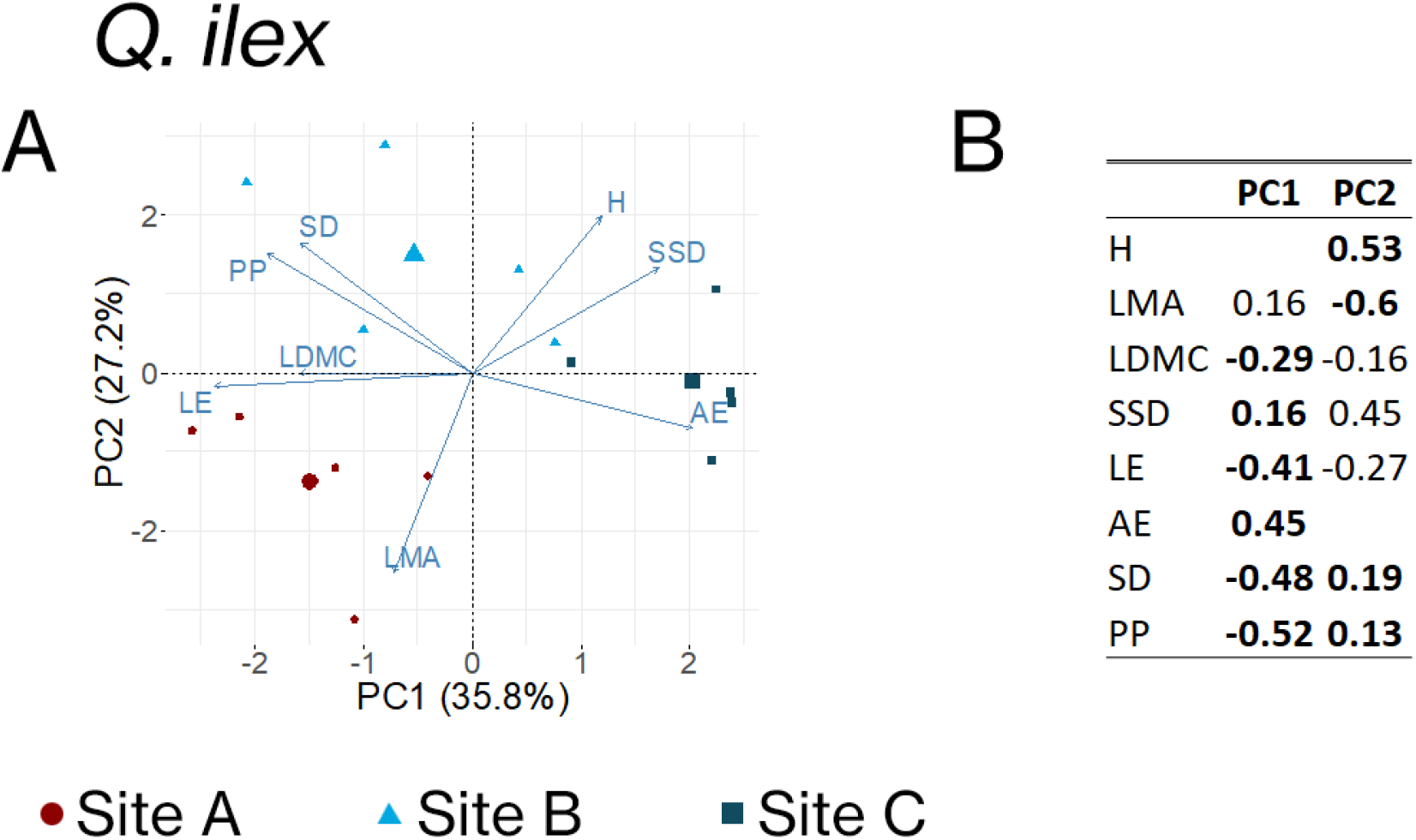
Principal Component Analysis (PCA) of trait variation for Q. *ilex,* with a biplot **A** showing the first two principal components and a table **B** showing the eigenvalues of the first two principal components, calculated on Q. *ilex* data. H = plant height; LMA = leaf mass area; LDMC = leaf dry matter content; SSD = stem specific density; LE = shoot length growth efficiency; AE = abaxial leaf epidermis thickness; SD = stomatal density; PP = ratio between palisade and patchy parenchyma thickness. Significant correlations between trait values and principal components are reported in bold in **B** (Pearson correlation test, p < 0.05).

TPD analysis clearly highlighted the differences between the distibutions of sites A and C for PC1 (Fig. 4A) and sites A and B for PC2 (Fig. 4B), with the third site always showing distributions similar to site A but slightly shifted towards the other site. Dissimilarity matrixes for the TPD are presented in Table 1.

**Figure 4.**
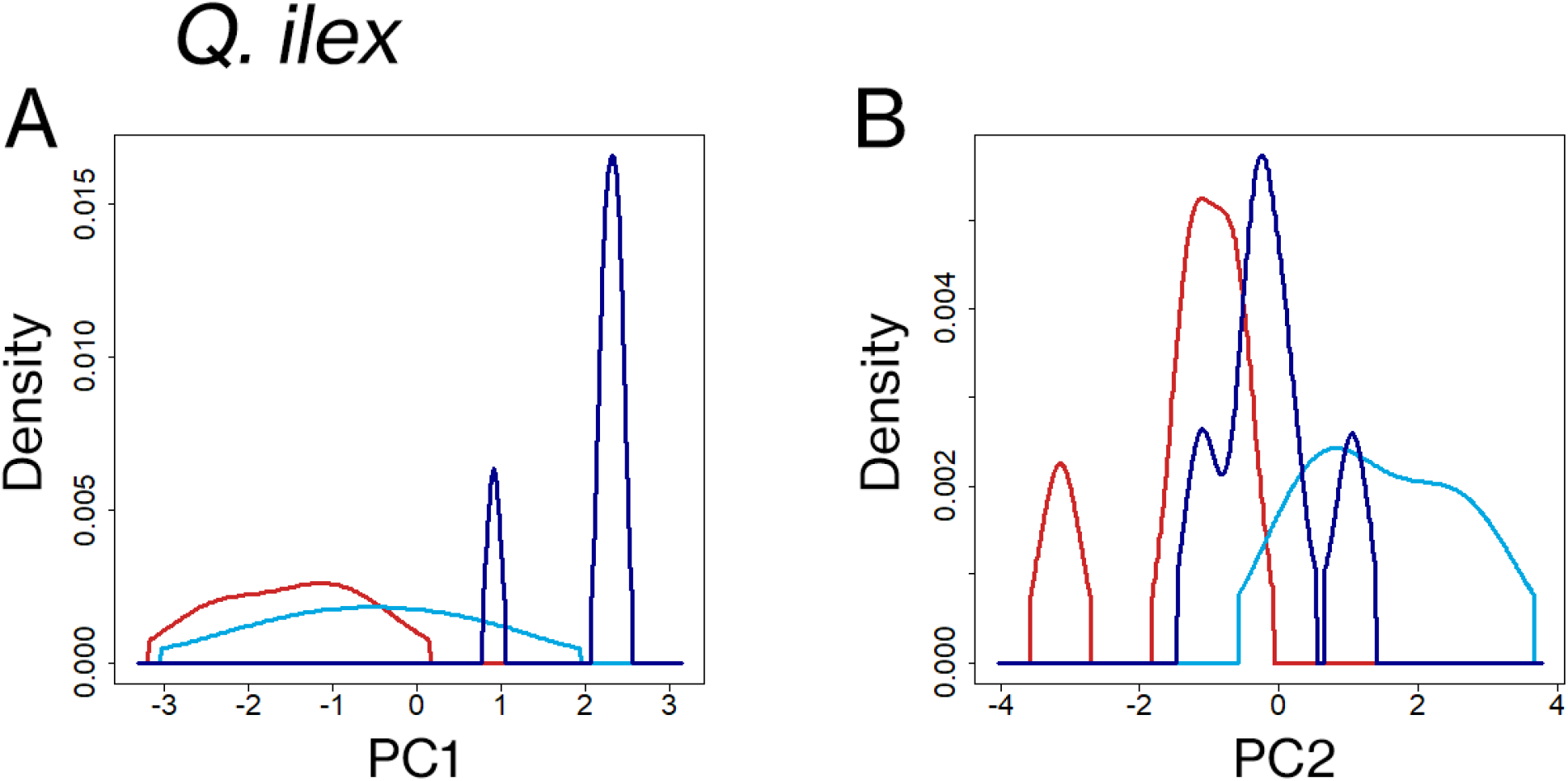
Trait Probability Density (TPD) distributions for the first two principal components of trait variation for Q. *ilex,* divided in **A** PC1 and **B** PC2. The distribution of plant functional traits from site A is represented in red, the distribution from site B is represented in light blue and the distribution from site C is represented in dark blue.

**Table 1.**
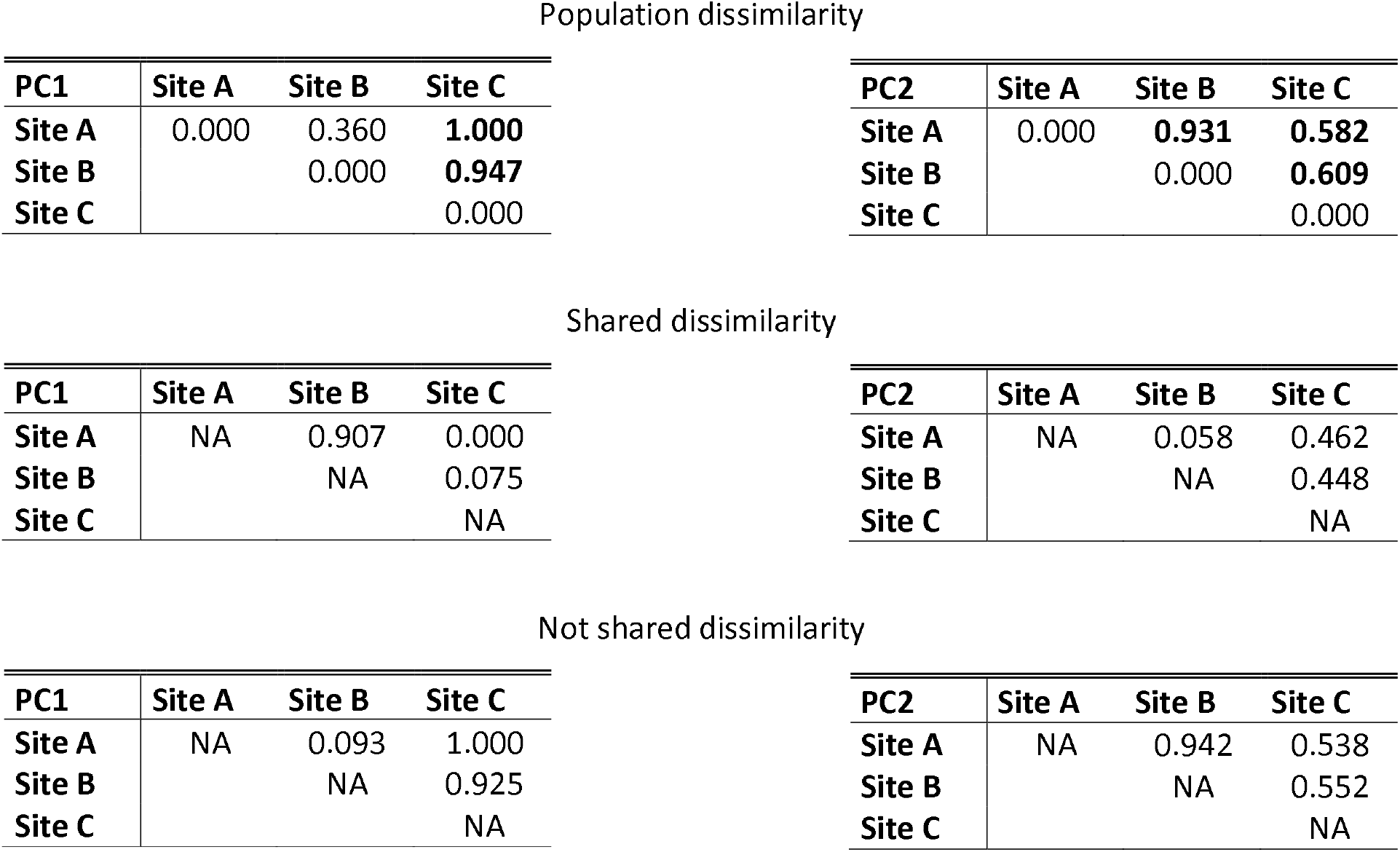
Dissimilarity matrixes of the first two principal components, calculated on Q. *ilex* data. Site A = Castel Fusano (RM) (3 m asl); Site B = La Farnesiana (RM) (150 m asl); Site C = Monte Catillo in Tivoli (RM) (430 m asl). Significant differences between sites are reported in bold (Wilcoxon-Mann-Whitney U test, p < 0.05).

Population dissimilarity
| PC1 | Site A | Site B | Site C |
| --- | --- | --- | --- |
| Site A | 0.000 | 0.360 | **1.000** |
| Site B |  | 0.000 | **0.947** |
| Site C |  |  | 0.000 |

| PC2 | Site A | Site B | Site C |
| --- | --- | --- | --- |
| Site A | 0.000 | **0.931** | **0.582** |
| Site B |  | 0.000 | **0.609** |
| Site C |  |  | 0.000 |

Shared dissimilarity
| PC1 | Site A | Site B | Site C |
| --- | --- | --- | --- |
| Site A | NA | 0.907 | 0.000 |
| Site B |  | NA | 0.075 |
| Site C |  |  | NA |

| PC2 | Site A | Site B | Site C |
| --- | --- | --- | --- |
| Site A | NA | 0.058 | 0.462 |
| Site B |  | NA | 0.448 |
| Site C |  |  | NA |

Not shared dissimilarity
| PC1 | Site A | Site B | Site C |
| --- | --- | --- | --- |
| Site A | NA | 0.093 | 1.000 |
| Site B |  | NA | 0.925 |
| Site C |  |  | NA |

**Table 1.** Dissimilarity matrixes of the first two principal components, calculated on *Q.
| PC2 | Site A | Site B | Site C |
| --- | --- | --- | --- |
| Site A | NA | 0.942 | 0.538 |
| Site B |  | NA | 0.552 |
| Site C |  |  | NA |

Population dissimilarity

Wilcoxon-Mann-Whitney U test (Table 1) confirmed the significant differences (p < 0.05) highlighted by the TPD between site C and the other ones on the first axis and between all the sites on the second axis, where H and LMA had greater influence.

### *Cistus salviifolius* L

The first two axes of *C. salviifolius* PCA (Fig. 5A) accounted for 54.8% of the total variance (31.3% by PC1 and 23.5% by PC2). PC1 resulted significantly related (p < 0.05) to SSD and inversely to H, LE and SD. PC2 was inversely related to H, LMA and LDMC, with a strong negative influence of the latter on the component (Fig. 5B). The analysis didn’t highlight marked differences across sites for this species.

**Figure 5.**
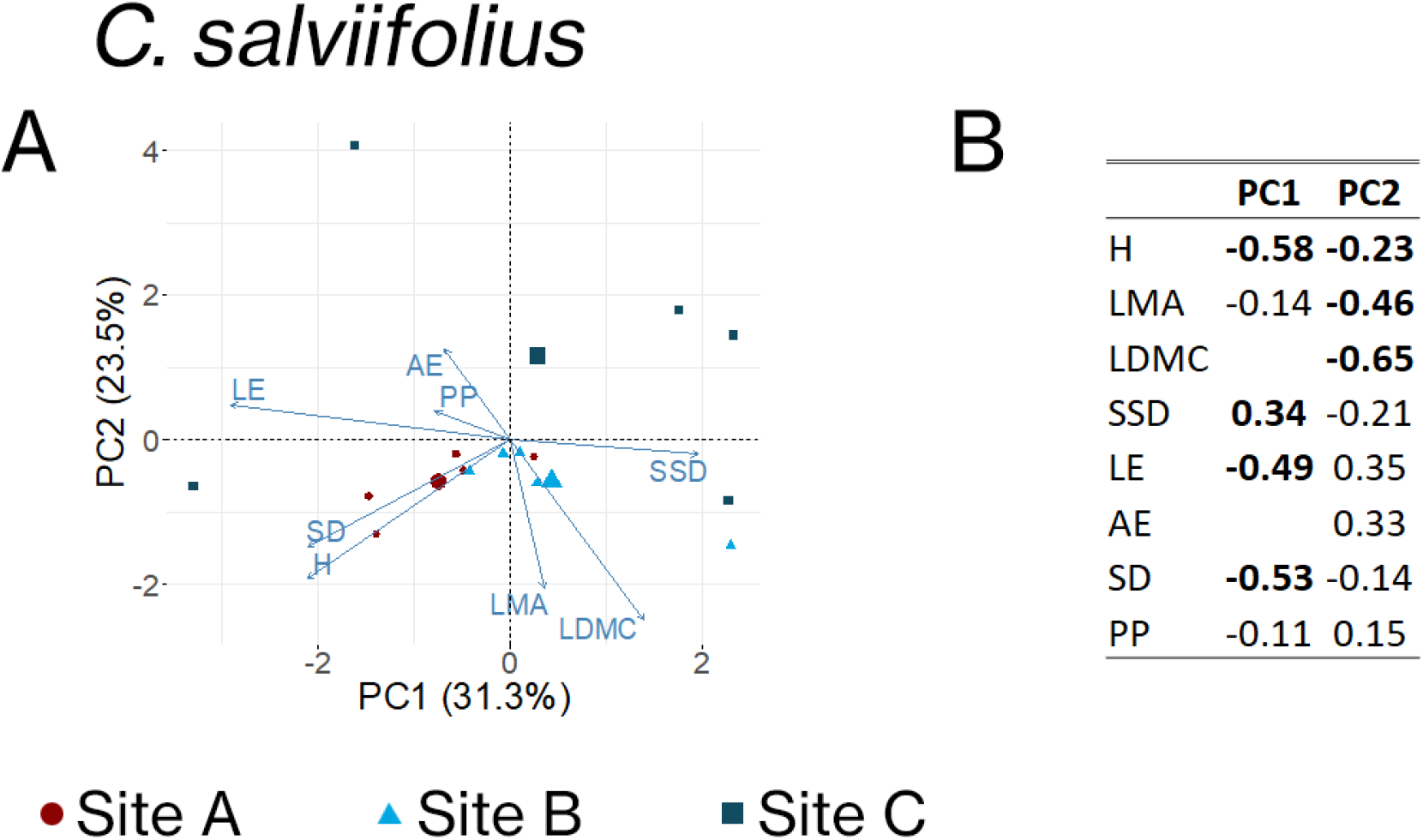
Principal Component Analysis (PCA) of trait variation for *C. salviifolius,* with a biplot **A** showing the first two principal components and a table **B** showing the eigenvalues of the first two principal components, calculated on *C. salviifolius* data. H = plant height; LMA = leaf mass area; LDMC = leaf dry matter content; SSD = stem specific density; LE = shoot length growth efficiency; AE = abaxial leaf epidermis thickness; SD = stomatal density; PP = ratio between palisade and patchy parenchyma thickness. Significant correlations between trait values and principal components are reported in bold in **B** (Pearson correlation test, p < 0.05).

TPD analysis showed an important overlap of sites distributions for both the first two axes of the PCA (Fig. 6), in particular among sites A and B, which also shared the shape of the distribution, while site C presented a flatter distribution.

**Figure 6.**
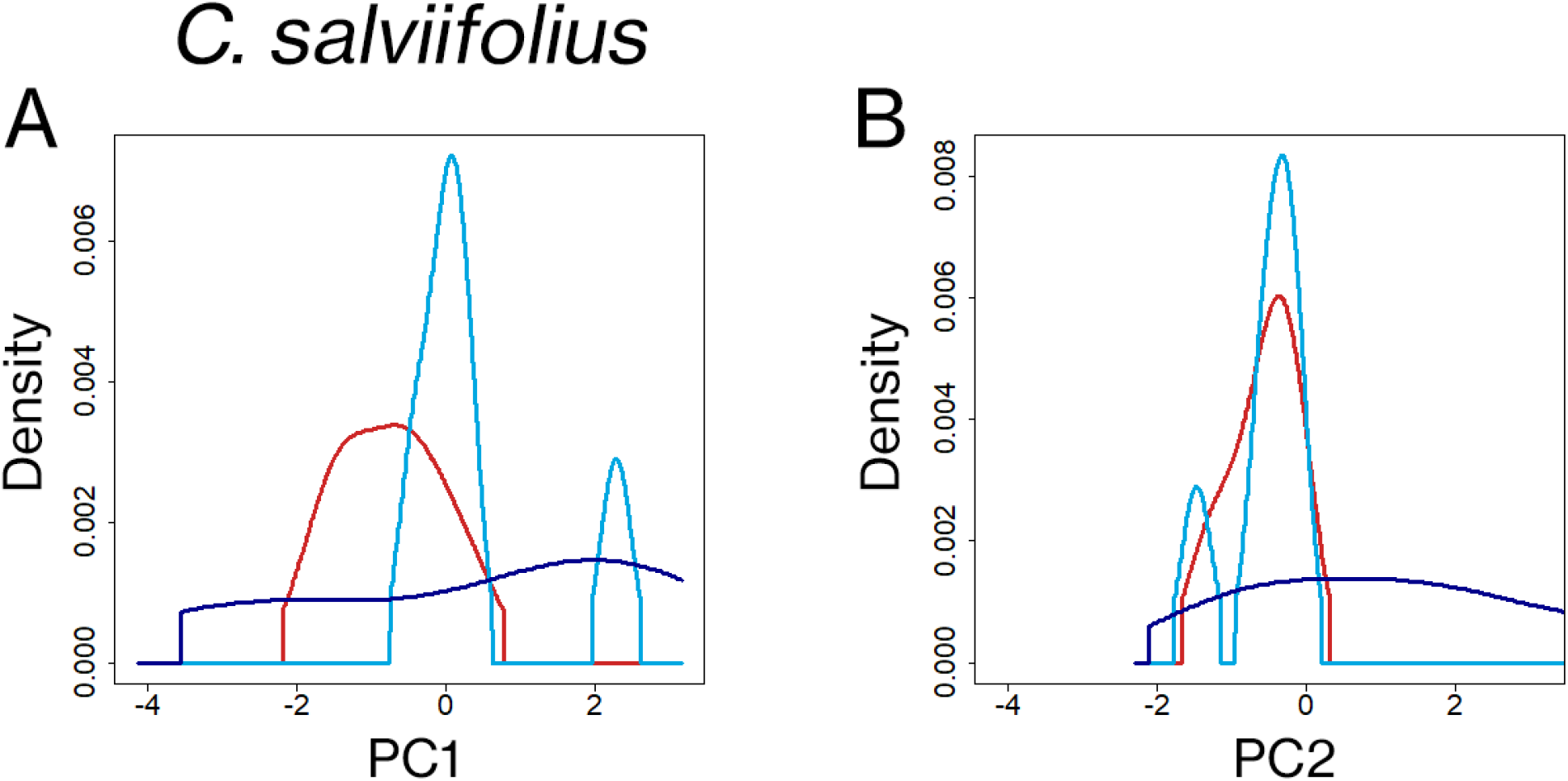
Trait Probability Density (TPD) distributions for the first two principal components of trait variation for *C. salviifolius*, divided in **A** PC1 and **B** PC2. The distribution of plant functional traits from site A is represented in red, the distribution from site B is represented in light blue and the distribution from site C is represented in dark blue.

Wilcoxon-Mann-Whitney U test (Table 2) confirmed the absence of significant differences between sites on both axes for this species.

**Table 2.**
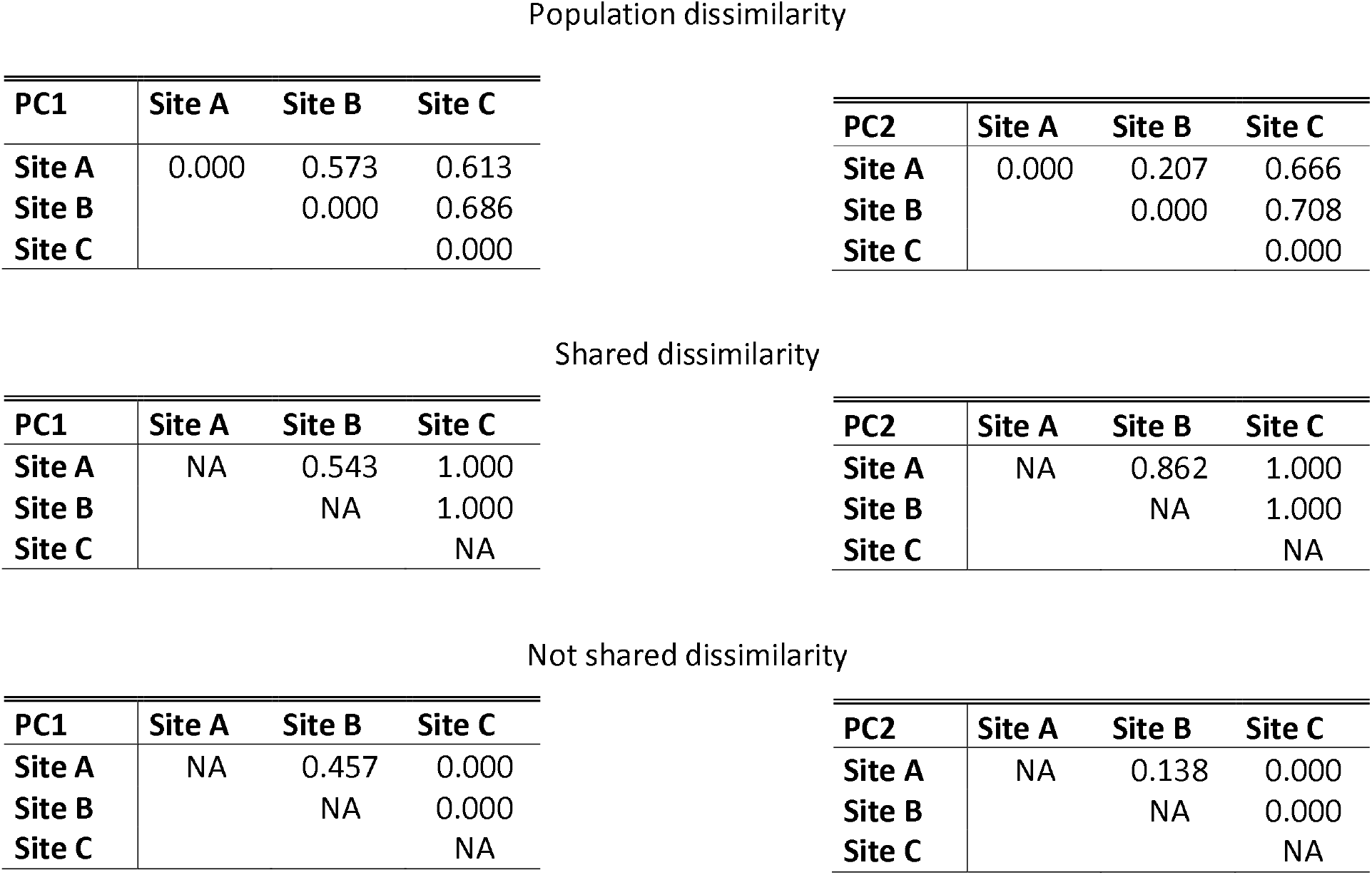
Dissimilarity matrixes of the first two principal components, calculated on *C. salviifolius* data. Site A = Castel Fusano (RM) (3 m asl); Site B = La Farnesiana (RM) (150 m asl); Site C = Monte Catillo in Tivoli (RM) (430 m asl). Significant differences between sites are reported in bold (Wilcoxon-Mann-Whitney U test, p < 0.05).

| PC1 | Site A | Site B | Site C |
| --- | --- | --- | --- |
| Site A | 0.000 | 0.573 | 0.613 |
| Site B |  | 0.000 | 0.686 |
| Site C |  |  | 0.000 |

| PC2 | Site A | Site B | Site C |
| --- | --- | --- | --- |
| Site A | 0.000 | 0.207 | 0.666 |
| Site B |  | 0.000 | 0.708 |
| Site C |  |  | 0.000 |

Shared dissimilarity
| PC1 | Site A | Site B | Site C |
| --- | --- | --- | --- |
| Site A | NA | 0.543 | 1.000 |
| Site B |  | NA | 1.000 |
| Site C |  |  | NA |

| PC2 | Site A | Site B | Site C |
| --- | --- | --- | --- |
| Site A | NA | 0.862 | 1.000 |
| Site B |  | NA | 1.000 |
| Site C |  |  | NA |

Not shared dissimilarity
| PC1 | Site A | Site B | Site C |
| --- | --- | --- | --- |
| Site A | NA | 0.457 | 0.000 |
| Site B |  | NA | 0.000 |
| Site C |  |  | NA |

**Table 2.** Dissimilarity matrixes of the first two principal components, calculated on *C.
| PC2 | Site A | Site B | Site C |
| --- | --- | --- | --- |
| Site A | NA | 0.138 | 0.000 |
| Site B |  | NA | 0.000 |
| Site C |  |  | NA |

### *Phyllirea latifolia* L

PCA carried on *P. latifolia* (Fig. 7A) data extracted two main axes explaining 81.7% of the total variance (60.6% by PC1 and 21.1% by PC2). PC1 was positively correlated (p < 0.05) with LMA, LE and AE, and negatively with SSD, SD, and PP. PC2 was positively correlated with H and negatively with LDMC (Fig. 7B). Sites separation along the axes was well defined. Site A showed higher values of LMA, LE and AE than sites B and C.

**Figure 7.**
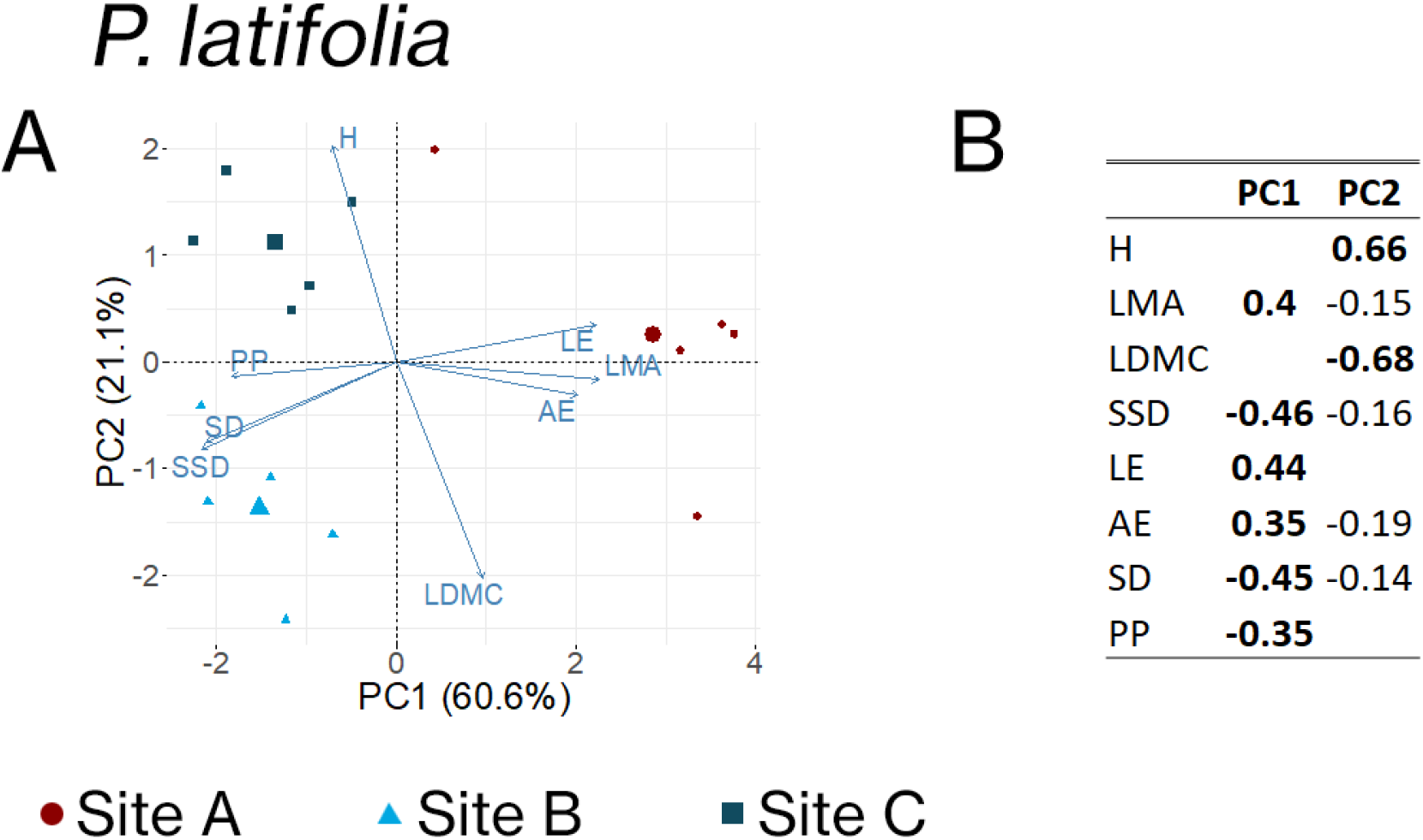
Principal Component Analysis (PCA) of trait variation for *P. latifolia,* with a biplot **A** showing the first two principal components and a table **B** showing the eigenvalues of the first two principal components, calculated on *P. latifolia* data. H = plant height; LMA = leaf mass area; LDMC = leaf dry matter content; SSD = stem specific density; LE = shoot length growth efficiency; AE = abaxial leaf epidermis thickness; SD = stomatal density; PP = ratio between palisade and patchy parenchyma thickness. Significant correlations between trait values and principal components are reported in bold in **B** (Pearson correlation test, p < 0.05).

TPD analysis displayed a strong difference between site A and the other ones and an overlap of distributions of sites B and C on PC1 (Fig. 8A). On the contrary,marked differences between the distributions of sites B and C were shown on PC2 (Fig. 8B).

**Figure 8.**
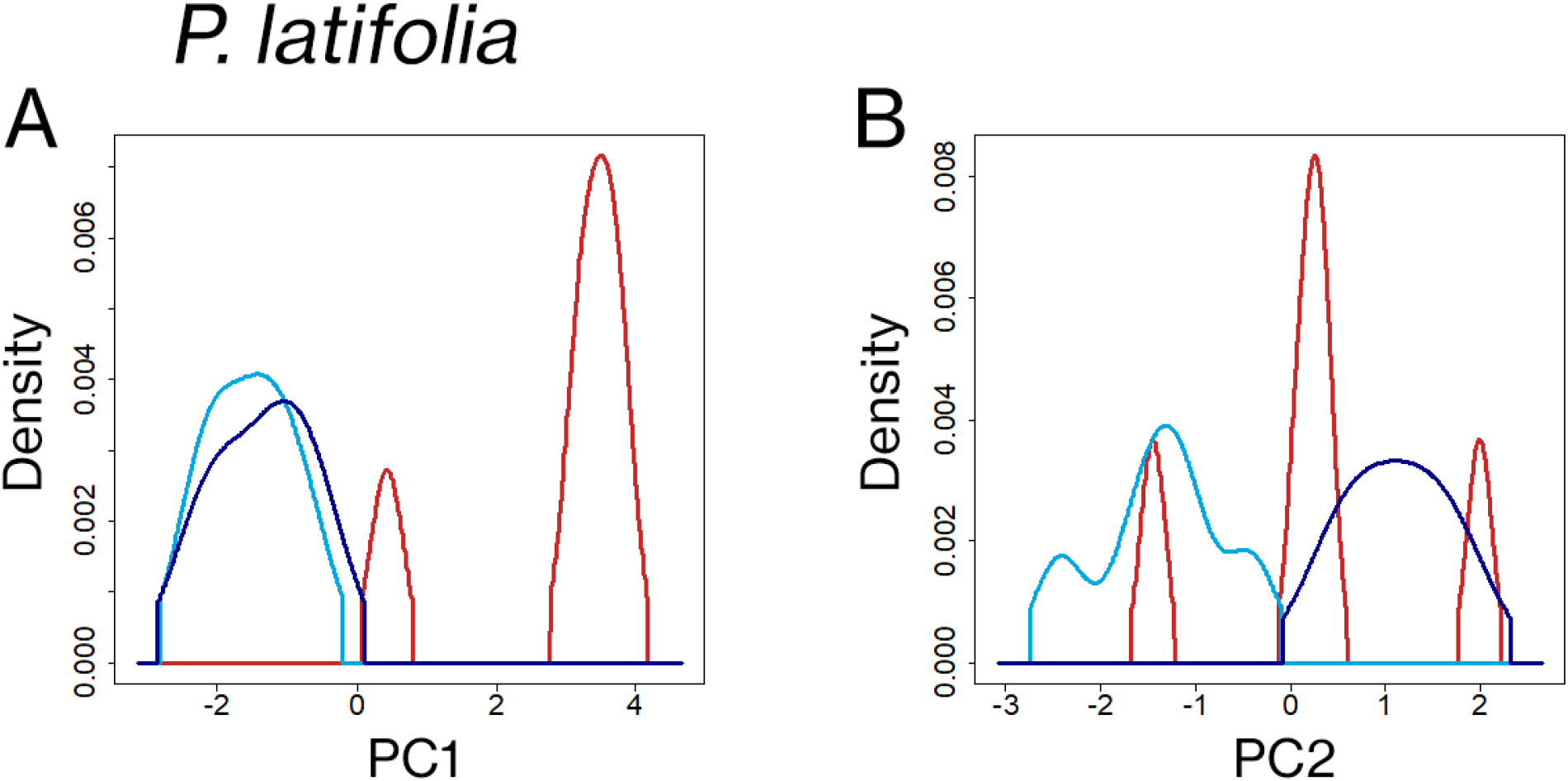
Trait Probability Density (TPD) distributions for the first two principal components of trait variation for *P. latifolia,* divided in **A** PC1 and **B** PC2. The distribution of plant functional traits from site A is represented in red, the distribution from site B is represented in light blue and the distribution from site C is represented in dark blue.

In agreement with TPD, Wilcoxon-Mann-Whitney U test (Table 3) detected significant differences (p < 0.05) between site A and the other ones on the first axis and between sites B and C on the second axis.

**Table 3.**
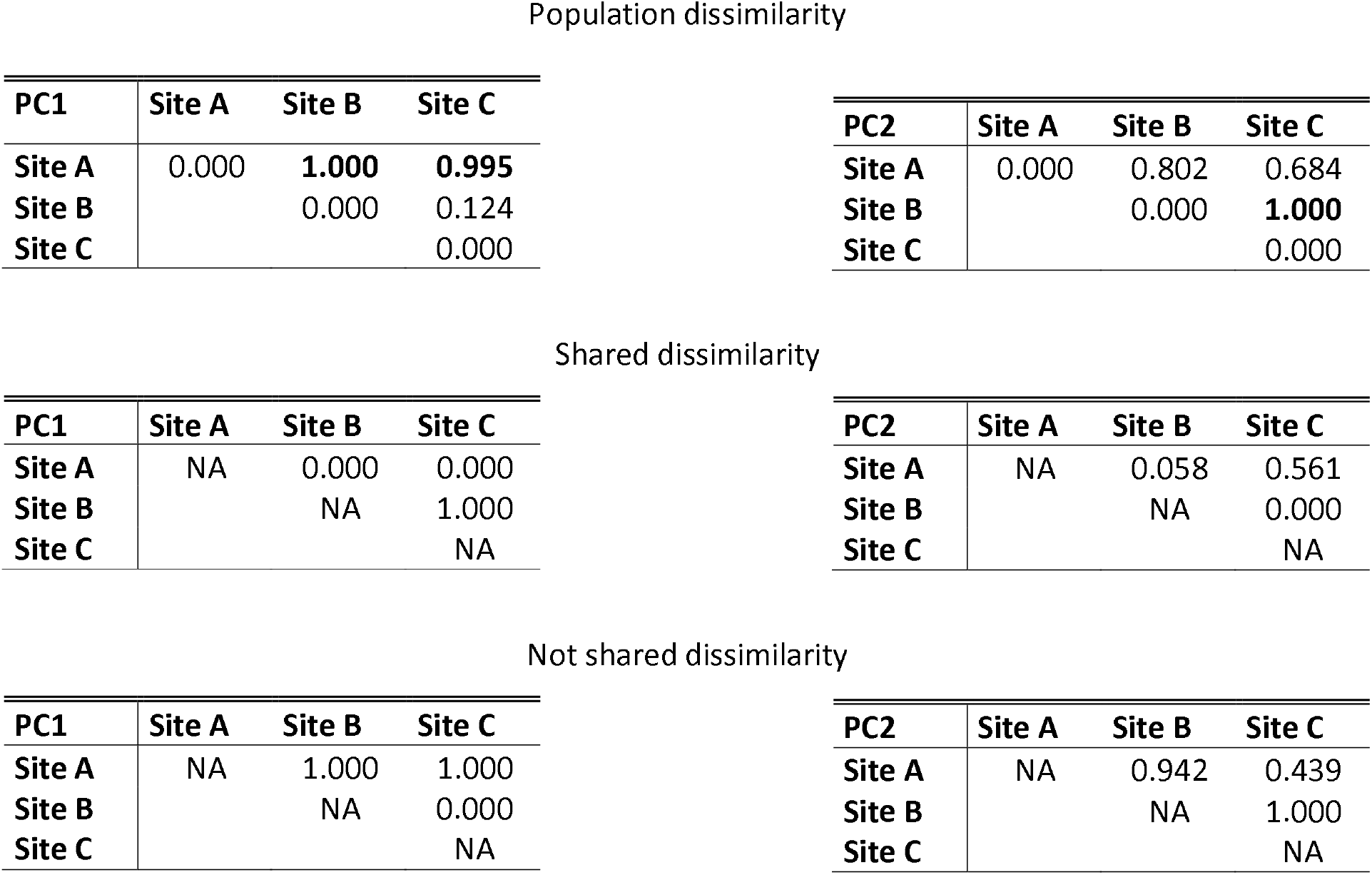
Dissimilarity matrixes of the first two principal components, calculated on *P. latifolia* data. Site A = Castel Fusano (RM) (3 m asl); Site B = La Farnesiana (RM) (150 m asl); Site C = Monte Catillo in Tivoli (RM) (430 m asl). Significant differences between sites are reported in bold (Wilcoxon-Mann-Whitney U test, p < 0.05).

| PC1 | Site A | Site B | Site C |
| --- | --- | --- | --- |
| Site A | 0.000 | **1.000** | **0.995** |
| Site B |  | 0.000 | 0.124 |
| Site C |  |  | 0.000 |

| PC2 | Site A | Site B | Site C |
| --- | --- | --- | --- |
| Site A | 0.000 | 0.802 | 0.684 |
| Site B |  | 0.000 | **1.000** |
| Site C |  |  | 0.000 |

Shared dissimilarity
| PC1 | Site A | Site B | Site C |
| --- | --- | --- | --- |
| Site A | NA | 0.000 | 0.000 |
| Site B |  | NA | 1.000 |
| Site C |  |  | NA |

| PC2 | Site A | Site B | Site C |
| --- | --- | --- | --- |
| Site A | NA | 0.058 | 0.561 |
| Site B |  | NA | 0.000 |
| Site C |  |  | NA |

Not shared dissimilarity
| PC1 | Site A | Site B | Site C |
| --- | --- | --- | --- |
| Site A | NA | 1.000 | 1.000 |
| Site B |  | NA | 0.000 |
| Site C |  |  | NA |

**Table 3.** Dissimilarity matrixes of the first two principal components, calculated on *P.
| PC2 | Site A | Site B | Site C |
| --- | --- | --- | --- |
| Site A | NA | 0.942 | 0.439 |
| Site B |  | NA | 1.000 |
| Site C |  |  | NA |

### *Pistacia lentiscus* L

PCA on *P. lentiscus* (Fig. 9A) data returned two axes of major variation accounting for the 37.2% (PC1) and 26.2% (PC2) of the total variance, for an amount of 63.4%. The first axis resulted significantly related (p < 0.05) to AE and inversely to LMA, LDMC and PP. The second axis was significantly related (p < 0.05) to LE and inversely to SSD and SD (Fig. 9B). PCA results underlined the difference between site C and the other sites, mostly as regards LMA, LDMC, H, AE and PP.

**Figure 9.**
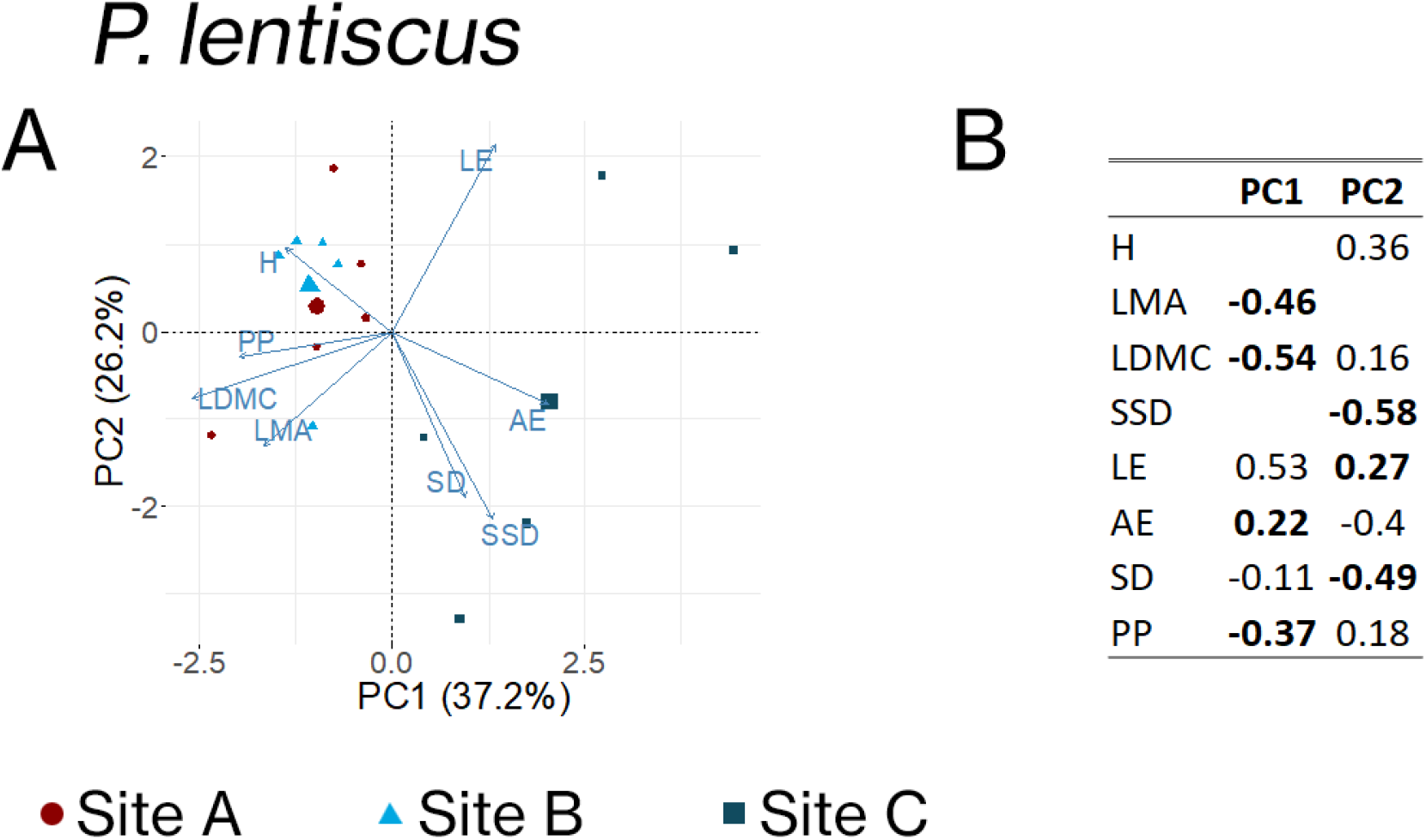
Principal Component Analysis (PCA) of trait variation for *P. lentiscus,* with a biplot **A** showing the first two principal components and a table **B** showing the eigenvalues of the first two principal components, calculated on *P. lentiscus* data. H = plant height; LMA = leaf mass area; LDMC = leaf dry matter content; SSD = stem specific density; LE = shoot length growth efficiency; AE = abaxial leaf epidermis thickness; SD = stomatal density; PP = ratio between palisade and patchy parenchyma thickness. Significant correlations between trait values and principal components are reported in bold in **B** (Pearson correlation test, p < 0.05).

TPD analysis displayed differences between site C and the other sites, with a partial overlap of sites A and B distributions on PC1 (Fig. 10A), while along PC2 the distributions did not show great differences (Fig. 10B).

**Figure 10.**
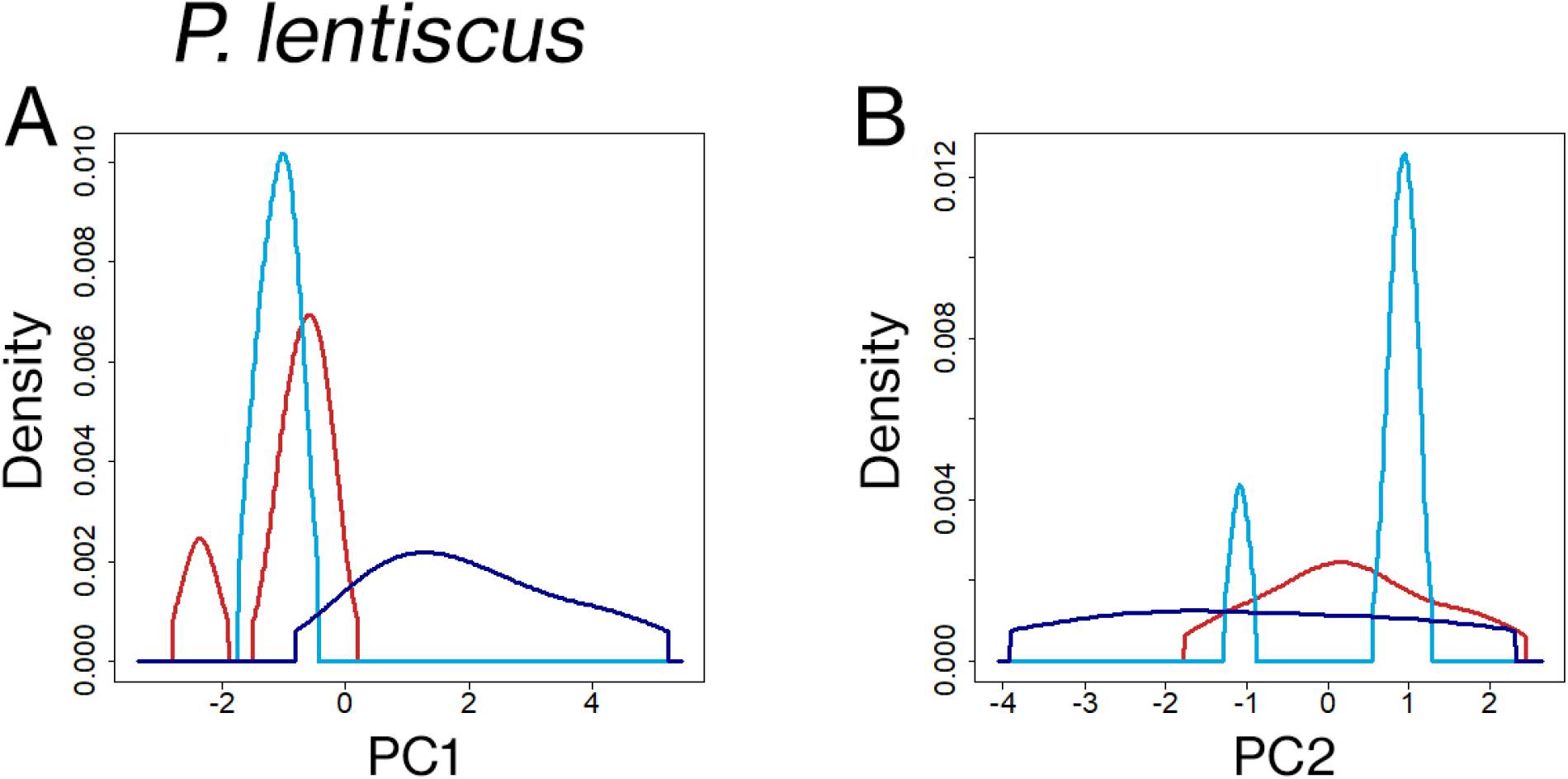
Trait Probability Density (TPD) distributions for the first two principal components of trait variation for *P. lentiscus,* divided in **A** PC1 and **B** PC2. The distribution of plant functional traits from site A is represented in red, the distribution from site B is represented in light blue and the distribution from site C is represented in dark blue.

In agreement with TPD analysis, Wilcoxon-Mann-Whitney U test (Table 4) reported significant differences (p < 0.05) between site C and the other ones on the first axis and no significative differences between sites on the second axis.

**Table 4.**
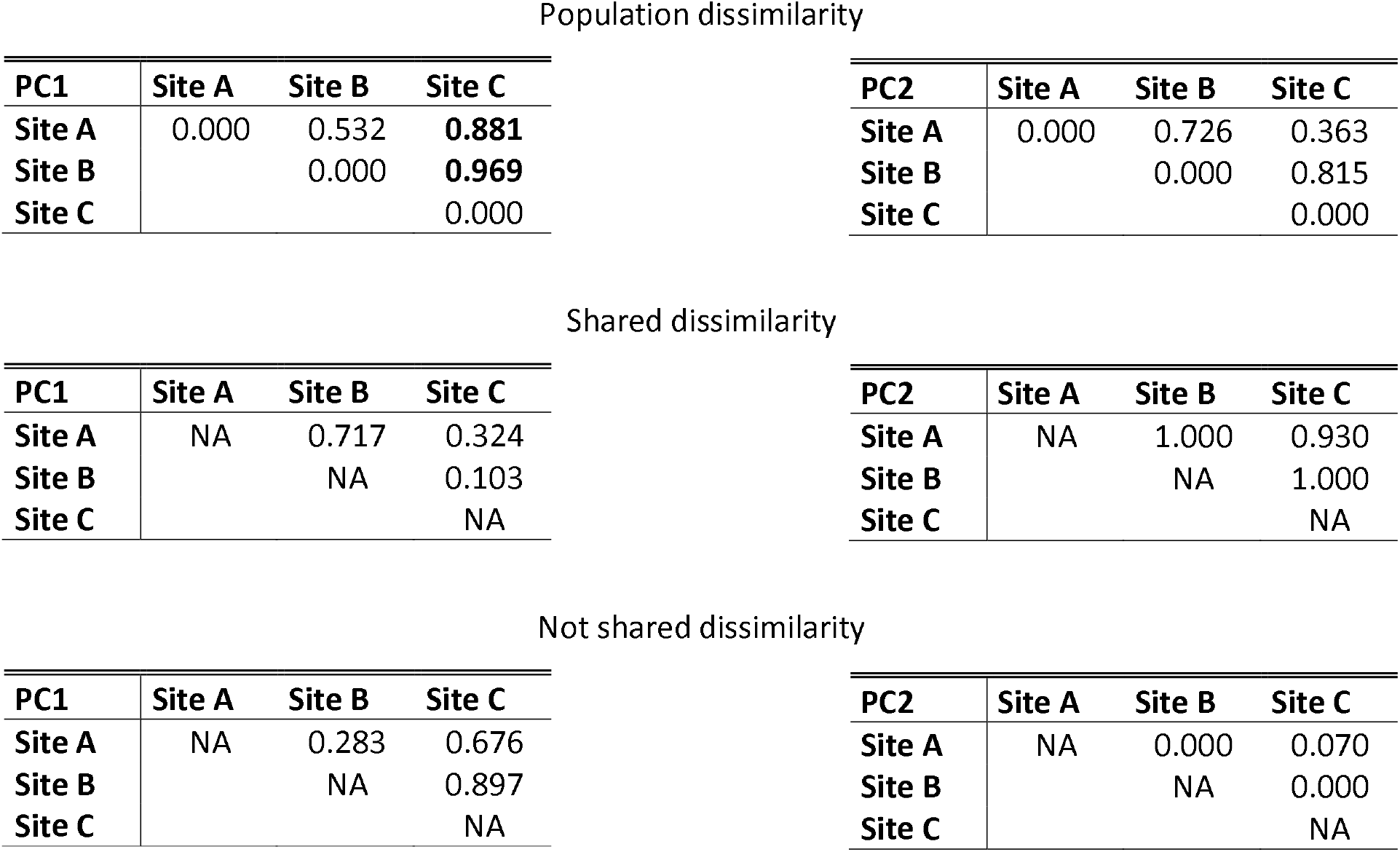
Dissimilarity matrixes of the first two principal components, calculated on *P. lentiscus* data. Site A = Castel Fusano (RM) (3 m asl); Site B = La Farnesiana (RM) (150 m asl); Site C = Monte Catillo in Tivoli (RM) (430 m asl). Significant differences between sites are reported in bold (Wilcoxon-Mann-Whitney U test, p < 0.05).

| PC1 | Site A | Site B | Site C |
| --- | --- | --- | --- |
| Site A | 0.000 | 0.532 | **0.881** |
| Site B |  | 0.000 | **0.969** |
| Site C |  |  | 0.000 |

| PC2 | Site A | Site B | Site C |
| --- | --- | --- | --- |
| Site A | 0.000 | 0.726 | 0.363 |
| Site B |  | 0.000 | 0.815 |
| Site C |  |  | 0.000 |

Shared dissimilarity
| PC1 | Site A | Site B | Site C |
| --- | --- | --- | --- |
| Site A | NA | 0.717 | 0.324 |
| Site B |  | NA | 0.103 |
| Site C |  |  | NA |

| PC2 | Site A | Site B | Site C |
| --- | --- | --- | --- |
| Site A | NA | 1.000 | 0.930 |
| Site B |  | NA | 1.000 |
| Site C |  |  | NA |

Not shared dissimilarity
| PC1 | Site A | Site B | Site C |
| --- | --- | --- | --- |
| Site A | NA | 0.283 | 0.676 |
| Site B |  | NA | 0.897 |
| Site C |  |  | NA |

**Table 4.** Dissimilarity matrixes of the first two principal components, calculated on *P.
| PC2 | Site A | Site B | Site C |
| --- | --- | --- | --- |
| Site A | NA | 0.000 | 0.070 |
| Site B |  | NA | 0.000 |
| Site C |  |  | NA |

## Discussion

In this study, we present evidence for significant differences in multiple PFTs among populations of several species compared at a local scale. Understanding plant’s potential for adaptation to climate is crucial, given the soon predicted increase of drought intensity and duration in the Mediterranean region (Diffenbaugh *et al*., 2007; Gao and Giorgi, 2008). Predicting the way species are going to respond to aridity is a complex task, which must involve an assessment of ITV to gain precision. This study aims at understanding the ITV of multiple PFTs from selected Mediterranean plant species, revealing insights into how local variations in climate drive differences between natural populations.

Q. *ilex* and *P. latifolia* present the strongest variability, while *P. lentiscus* appears slightly less variable and *C. salviifolius* fails to show significant differences among most PFTs in the studied sites. Analysed traits often show a significant amount of ITV although the overall variability is, as reasonably expected, mainly driven by BTV. The amount of observed ITV (0.113) may seem modest but is astounding when compared to the estimated global amount of ITV (0.25, Siefert *et al*., 2015), given the populations we studied only cover a small fraction of their whole species distributions. The region we chose for this study conveniently offers a wide climatic space in small spatial area, proving to be ideal to demonstrate the primary role of climate over locality in plant adaptation as suggested by Hufford and Mazer (2003).

Observed variations are consistent with PFTs adaptations to aridity previously reported in literature (Crescente *et al*., 2002; Wright *et al*., 2005; Ivanov *et al*., 2008; Martínez-Cabrera *et al*., 2009; Carlson *et al*., 2016; Guo *et al*., 2017; Nunes *et al*., 2017; Dörken *et al*., 2020; Anderegg *et al*., 2021): individuals hailing from populations thriving in drier sites generally show higher amounts of LMA, LDMC, LE and palisade on spongy parenchyma ratio coupled with lower plant height and specific stem density. Some functional traits show more erratic responses to drought among species, such as abaxial leaf epidermis thickness and stomatal density which tend to either increase or decrease with climate variations.

However, it is important to step further from single PFTs comparisons and consider population changes between a multivariate functional space to better understand how coordinated changes might lead to alternative strategies to resist drought (Blumenthal *et al*., 2020; Rodríguez-Alarcón *et al*., 2022). Water optimization can be achieved through multiple strategies, sometimes involving the adoption of apparently discordant responses of the same PFTs. Such is the case of the erratic response we observed for SD, which can accomplish its goal of optimizing fitness to aridity with either a decrease (aimed at reducing water losses) or an increase (aimed at cooling down the leaf and accurately controlling gas exchanges). Some have even argued that functional traits like SD do not show a tight relationship with water loss, concluding that direct relationships between traits and physiological responses are context-dependent (Carlson *et al*., 2016). It is indeed the context that allows to comprehend functional strategies, and PFTs need to be examined in syndromes to thoroughly understand their strategic adoption.

The adoption of the observed PFTs syndromes allows plants living in dry sites to adopt a conservative strategy, aimed at increasing leaves’ ability to withstand periods of more intense stress at the expense of a higher physiological cost for the plant. By trading off leaf production costs for thicker, more dense leaves, plants can optimize their survivability during drought, investing in leaves that can withstand higher stress and minimize water losses. The observed lower plant height in dry sites may reduce the risk of cavitation under increased water stress (Nunes *et al*., 2017), although it’s still possible that closer proximity to the sea (< 2 Km) could also induce lower plant height as a response to winds carrying salt spray (Du and Hesp, 2020). Plants hailing from the dry site also benefit from increased shoot length growth efficiency, probably driven by an earlier start of the vegetative season due to warmer temperatures.

Contrarily to our expectation, plants hailing from the intermediate site do not usually show intermediate trait values. Instead, they often adopt traits similar to either the dry site or the wet site, with a generally inconsistent pattern. Such a behaviour could derive from the fact that natural selection does not necessarily drive the adoption of changes when unoptimized traits are still fairly performing. An optimized strategy could only be necessary after a critical ecological threshold is reached, allowing for the preservation of unoptimized PFTs in intermediate conditions (Zhang *et al*., 2021; Virta and Teittinen, 2022). A possible solution to this riddle could reside in the phylogenetic history of our species. If we consider the distribution of trait syndromes over the major axes of variation, we notice that site B is usually similar to site A except for *P. latifolia,* for which the intermediate site mostly adopts PFTs similar to the wet site. While *C. salviifolius* and possibly Q. *ilex* evolved in Mediterranean conditions (Correia and Catarino, 1994; Blondel and Aronson, 1999; Martín-Sánchez *et al*., 2022) and *P. lentiscus* evolved in the semi-arid steppes of Central Asia (Blondel and Aronson, 1999), *P. latifolia* evolved in an arid Afro-Tropical context (Quézel, 1985). This means that *P. latifolia* could have lacked the pre-adaptation to be competitive with the amount of rain present at site C, inducing the need to adopt specific adaptations to populate that area. Likewise, the other species may have needed to adopt specific adaptations to thrive at site a despite the higher temperatures and lower precipitations, thus explaining the different PFTs syndromes adopted by each population.

Regardless of when the transition happened, we can still appreciate how climate variations are able to significantly affect ITV on a local scale. The adoption of PFTs widely regarded as adaptations to aridity is indeed a clue that climate could be the main driver for the observed changes in our data.

Plants responses to climate on a local scale are also species-specific. The evergreens generally show a higher response to climate than the semi-deciduous shrub *C. salvifolius,* and this could partially be explained by the ability of this species to optimise its seasonal climatic fitness by switching from summer to winter leaves (Grant *et al*., 2014). However, this does not explain why different populations sporting summer leaves do not further optimize their strategies to local differences in climate.

The answer may lie in the ecological strategy chosen by this species. As a pioneer species (Grant *et al*., 2014), *C. salviifolius* can thrive in a lacking competitor’s environment thanks to a set of more stable and versatile, although less specialised, functional traits. Versatile traits allow to rapidly colonise pristine habitats whereas specialised plants would need time to adapt, a resource that is essential when initiative determines success. The species that showed a more evident response to small-scale climate gradients are indeed those that are more specialised and competitive in advanced seral stages and that need to develop specific adaptations, as hypothesised by Reich (2014). Recent literature seems to support this explanation: in a study on the role of ITV during succession, Chai *et al.* (2019) found a considerable effect of ITV only at the latest climax communities, transitioning from the stochastic pattern of the early communities. On the other hand, the arboreal *Eremanthus erythropappus* (DC.) MacLeish is a dominant pioneer species with conservative water use, that was found to show significant ITV in forest-savannah ecotones (Silva *et al*., 2019). This might also imply that structural differences (i.e., herbaceous vs arboreal) are also important factors in determining ITV among species.

Another issue that arises is that overall, our results highlight the presence of ITV but cannot resolve the origin of the observed variability. To further consider this possibility and thoroughly comprehend species adaptability, it is crucial to understand whether the observed variations derive from morphological plasticity (i.e., the ability of a single genotype to adopt different phenotypes in response to the environment) or genetic adaptation, the two main sources of ITV (de Bello *et al*., 2021).

Resolving this ambiguity could lead to interesting implications for conservation biology as well: in presence of genetically divergent ecotypes, it would be crucial to consider populations, not species, as target entities to protect. Considering this, seeds from the populations presented in this study have already been collected with the aim of carrying out a common garden experiment. Combined with induced aridity, we aim to further investigate the ties between the data we observed and climate adaptation.

In conclusion, we suggest to use caution when assuming homogenous functional strategies among populations living close to each other. Environmental differences proved to be able to affect ITV at a local scale, pushing several plant species to adopt opportune strategies to cope with local stresses. The data we presented supports the idea that ITV behaves differently among species, implying that phylogenetic history, structural habit and even seral stages could be drivers of these changes. Given these premises, we invite to thoroughly consider whether it would be the case to shift the focus from species to populations when considering the possible responses of plants to a changing climate.

## ABBREVIATIONS

AE: Thickness of the Abaxial Leaf Epidermis
BTV: Between-species Variability
DM: Leaf Dry Mass
DM_S_: Shoot Leaf Dry Mass
H: Mean Plant height
ITV: Intraspecific Variability
LA: Leaf Area
LA_S_: Shoot Leaf Surface Area
LDMC: Leaf Dry Matter Content
LE: Shoot Length Growth Efficiency
LMA: Leaf Mass per Area
_Pal_PT: Palisade Parenchyma Thickness
_pat_PT: Patchy Parenchyma Thickness
PCA: Principal Components Analysis
PFTs: Plant Functional Traits
PP: Palisade over Parenchyma Thickness Ratio
SD: Stomatal Density
SDM_s_: Shoot Stem Dry Mass
SD_s_: Shoot Stem Diameter
SL_s_: Shoot Stem Length
SM: Leaf Water Saturated Mass
SSD: Specific Stem Density
SV_s_: Shoot Stem Volume
TPD: Trait Probability Distribution Analysis
VPA: Variance Partition Analysis

## SUPPLEMENTARY DATA

The following supplementary data are available at JXB online.

Table S1. Chemical-physical soil analysis data.

Table S2. Functional traits data, divided for each species.

## ACKNOWLEDGEMENTS

We thank Dr. Giuseppe Fabrini for his help on identifying species and Dr. Giacomo Puglielli for his constructive suggestions on statistical analyses. We also warmly thank Alessia Massimi and Maria Sacchettoni for their kind help with the fieldwork.

## AUTHOR CONTRIBUTIONS

Author Contributions: L.M.I. contributed to conceiving the idea, design the experiment and collecting field data. He also performed data analysis and wrote the manuscript. V.C. contributed on the collection of field data and writing the manuscript. L.V. conceived the idea, designed the experiment, and led the writing of the manuscript. All authors contributed critically to the final manuscript and gave their approval for publication.

## DATA AVAILABILITY

The data supporting the findings of this study are available from the corresponding author, Prof. Laura Varone, upon request.

## REFERENCES

Albert CH, Grassein F, Schurr FM, Vieilledent G, Violle C. 2011. When and how should intraspecific variability be considered in trait-based plant ecology? Perspectives in Plant Ecology, Evolution and Systematics 13, 217–225.

Anderegg LDL, Loy X, Markham IP, Elmer CM, Hovenden MJ, HilleRisLambers J, Mayfield MM. 2021. Aridity drives coordinated trait shifts but not decreased trait variance across the geographic range of eight Australian trees. New Phytologist 229, 1375–1387.

Basu S, Ramegowda V, Kumar A, Pereira A. 2016. Plant adaptation to drought stress. F1000Research 5.

de Bello F, Carmona CP, Dias AT, Götzenberger L, Moretti M, Berg MP. 2021. Handbook of trait-based ecology: from theory to R tools. Cambridge University Press.

de Bello F, Lavorel S, Albert CH, Thuiller W, Grigulis K, Dolezal J, Janeček Š, Lepš J. 2011. Quantifying the relevance of intraspecific trait variability for functional diversity. Methods in Ecology and Evolution 2, 163–174.

Blasi C. 1994. Fitoclimatologia del lazio. Univ. La Sapienza: Regione Lazio, Assessorato Agricoltura-Foreste, Caccia e Pesca, Usi Civici.

Blondel J, Aronson J. 1999. Biology and wildlife of the Mediterranean region. Oxford University Press, USA.

Blumenthal DM, Mueller KE, Kray JA, Ocheltree TW, Augustine DJ, Wilcox KR. 2020. Traits link drought resistance with herbivore defence and plant economics in semi-arid grasslands: The central roles of phenology and leaf dry matter content. Journal of Ecology 108, 2336–2351.

Carlson JE, Adams CA, Holsinger KE. 2016. Intraspecific variation in stomatal traits, leaf traits and physiology reflects adaptation along aridity gradients in a South African shrub. Annals of Botany 117, 195–207.

Carmona CP, Bello F, Mason NWH, Lepš J. 2019. Trait probability density (TPD): measuring functional diversity across scales based on TPD with R. Ecology 100, e02876.

Chabot BF, Chabot JF. 1977. Effects of light and temperature on leaf anatomy and photosynthesis in Fragaria vesca. Oecologia 26, 363–377.

Chai Y, Dang H, Yue M, et al. 2019. The role of intraspecific trait variability and soil properties in community assembly during forest secondary succession. Ecosphere 10, e02940.

Cornwell WK, Cornelissen JHC, Amatangelo K, et al. 2008. Plant species traits are the predominant control on litter decomposition rates within biomes worldwide. Ecology Letters 11, 1065–1071.

Correia O, Catarino F. 1994. Seasonal changes in soil-to-leaf resistance in *Cistus* sp. and *Pistacia lentiscus*. Acta oecologica (Montrouge) 15, 289–300.

Crescente MF, Gratani L, Larcher W. 2002. Shoot growth efficiency and production of *Quercus ilex* L. in different climates. Flora - Morphology, Distribution, Functional Ecology of Plants 197, 2–9.

Dalia Vecchia A, Bolpagni R. 2022. The importance of being petioled: leaf traits and resource-use strategies in Nuphar lutea. Hydrobiologia, 1–12.

Diffenbaugh NS, Pal JS, Giorgi F, Gao X. 2007. Heat stress intensification in the Mediterranean climate change hotspot. Geophysical Research Letters 34.

Dörken VM, Ladd PG, Parsons RF. 2020. Anatomical aspects of xeromorphy in arid-adapted plants of Australia. Australian Journal of Botany 68, 245–266.

Du J, Hesp PA. 2020. Salt Spray Distribution and Its Impact on Vegetation Zonation on Coastal Dunes: a Review. Estuaries and Coasts 43, 1885–1907.

Ferreira de Melo Junior JC, Torres Boeger MR. 2016. Leaf traits and plastic potential of plant species in a light-edaphic gradient from restinga in southern Brazil. Acta Biológica Colombiana 21, 51–62.

Fick SE, Hijmans RJ. 2017. WorldClim 2: new 1-km spatial resolution climate surfaces for global land areas. International Journal of Climatology 37, 4302–4315.

Funk JL, Larson JE, Ames GM, Butterfield BJ, Cavender-Bares J, Firn J, Laughlin DC, Sutton-Grier AE, Williams L, Wright J. 2017. Revisiting the Holy Grail: using plant functional traits to understand ecological processes. Biological Reviews 92, 1156–1173.

Gao X, Giorgi F. 2008. Increased aridity in the Mediterranean region under greenhouse gas forcing estimated from high resolution simulations with a regional climate model. Global and Planetary Change 62, 195–209.

Garcia MN, Hu J, Domingues TF, Groenendijk P, Oliveira RS, Costa FRC. 2022. Local hydrological gradients structure high intraspecific variability in plant hydraulic traits in two dominant central Amazonian tree species. Journal of Experimental Botany 73, 939–952.

Garrett KA, Zúñiga LN, Roncal E, Forbes GA, Mundt CC, Su Z, Nelson RJ. 2009. Intraspecific functional diversity in hosts and its effect on disease risk across a climatic gradient. Ecological Applications 19, 1868–1883.

Gaussen H, Bagnouls F. 1953. Saison sèche et indice xérothermique. Toulouse, França: Université de Toulouse, Facultei dês Sciences.

Grant OM, Tronina Ł, García-Plazaola JI, et al. 2014. Resilience of a semi-deciduous shrub, *Cistus salvifolius,* to severe summer drought and heat stress. Functional Plant Biology 42, 219–228.

Gratani L, Bombelli A. 2001. Differences in leaf traits among Mediterranean broad-leaved evergreen shrubs. JSTOR, 15–24.

Gratani L, Crescente MF. 1997. Phenology and leaf adaptive strategies of Mediterranean maquis plants. Ecologia Mediterranea 23, 11–19.

Gratani L, Varone L. 2004. Adaptive photosynthetic strategies of the Mediterranean maquis species according to their origin. Photosynthetica 42, 551–558.

Gratani L, Varone L, Crescente MF, Catoni R, Ricotta C, Puglielli G. 2018. Leaf thickness and density drive the responsiveness of photosynthesis to air temperature in Mediterranean species according to their leaf habitus. Journal of Arid Environments 150, 9–14.

Guo C, Ma L, Yuan S, Wang R. 2017. Morphological, physiological and anatomical traits of plant functional types in temperate grasslands along a large-scale aridity gradient in northeastern China. Scientific Reports 7, 40900.

Hall DO, Scurlock J, Bolhar-Nordenkampf H, Leegood RC, Long S. 2013. Photosynthesis and production in a changing environment: a field and laboratory manual. Springer Science & Business Media.

Harley PC, Tenhunen JD, Beyschlag W, Lange OL. 1987. Seasonal changes in net photosynthesis rates and photosynthetic capacity in leaves of *Cistus salvifolius,* a European mediterranean semi-deciduous shrub. Oecologia 74, 380–388.

Hufford KM, Mazer SJ. 2003. Plant ecotypes: genetic differentiation in the age of ecological restoration. Trends in Ecology & Evolution 18, 147–155.

Ivanov LA, Ronzhina DA, Ivanova LA. 2008. Changes in leaf characteristics as indicator of the alteration of functional types of steppe plants along the aridity gradient. Russian Journal of Plant Physiology 55, 301–307.

Jones >Jr JB. 1999. Soil analysis handbook of reference methods. CRC press.

Jung V, Albert CH, Violle C, Kunstler G, Loucougaray G, Spiegelberger T. 2014. Intraspecific trait variability mediates the response of subalpine grassland communities to extreme drought events. Journal of Ecology 102, 45–53.

Jung V, Violle C, Mondy C, Hoffmann L, Muller S. 2010. Intraspecific variability and trait-based community assembly. Journal of Ecology 98, 1134–1140.

Karger DN, Conrad O, Böhner J, Kawohl T, Kreft H, Soria-Auza RW, Zimmermann NE, Linder HP, Kessler M. 2017. Climatologies at high resolution for the earth’s land surface areas. Scientific Data 4, 170122.

King DA. 1997. Branch growth and biomass allocation in *Abies amabilis* saplings in contrasting light environments. Tree physiology 17, 251–258.

Kumordzi BB, Nilsson M-C, Gundale MJ, Wardle DA. 2014. Changes in local-scale intraspecific trait variability of dominant species across contrasting island ecosystems. Ecosphere 5, 1–17.

Larcher W. 2003. Physiological plant ecology: ecophysiology and stress physiology of functional groups. Springer Science & Business Media.

Lecerf A, Chauvet E. 2008. Intraspecific variability in leaf traits strongly affects alder leaf decomposition in a stream. Basic and Applied Ecology 9, 598–605.

Lionello P, Malanotte-Rizzoli P, Boscolo R. 2006. Mediterranean climate variability. Elsevier.

Maharjan SK, Sterck FJ, Dhakal BP, Makri M, Poorter L. 2021. Functional traits shape tree species distribution in the Himalayas. Journal of Ecology 109, 3818–3834.

Martinez-Cabrera HI, Jones CS, Espino S, Schenk HJ. 2009. Wood anatomy and wood density in shrubs: Responses to varying aridity along transcontinental transects. American Journal of Botany 96, 1388–1398.

Martín-Sánchez R, Peguero-Pina JJ, Alonso-Forn D, Ferrio JP, Sancho-Knapik D, Gil-Pelegrín E. 2022. Summer and winter can equally stress holm oak *(Quercus ilex* L.) in Mediterranean areas: A physiological view. Flora 290, 152058.

Moreira B, Tavsanoglu Ç, Pausas JG. 2012. Local versus regional intraspecific variability in regeneration traits. Oecologia 168, 671–677.

Nunes A, Köbel M, Pinho P, Matos P, Bello F de, Correia O, Branquinho C. 2017. Which plant traits respond to aridity? A critical step to assess functional diversity in Mediterranean drylands. Agricultural and Forest Meteorology 239, 176–184.

Oksanen J, Blanchet FG, Kindt R, Legendre P, Minchin PR, O’hara RB, Simpson GL, Solymos P, Stevens MHH, Wagner H. 2013. Package ‘vegan’. Community ecology package, version 2, 1–295.

Pérez-Harguindeguy N, Díaz S, Garnier E, et al. 2016. *Corrigendum* to: New handbook for standardised measurement of plant functional traits worldwide. Australian Journal of Botany 64, 715–716.

Pesoli P, Gratani L, Larcher W. 2003. Responses of *Quercus ilex* from different provenances to experimentally imposed water stress. Biologia plantarum 46, 577–581.

Puglielli G, Carmona CP., Varone L, Laanisto L, Ricotta C. 2022. Phenotypic dissimilarity index: Correcting for intra and interindividual variability when quantifying phenotypic variation. Ecology e3806.

Quezel P. 1985. Definition of the Mediterranean region and the origin of its flora. Geobotany.

Reich PB. 2014. The world-wide ‘fast–slow’ plant economics spectrum: a traits manifesto. Journal of Ecology 102, 275–301.

Rodríguez-Alarcón S, Tamme R, Carmona CP. 2022. Intraspecific trait changes in response to drought lead to trait convergence between—but not within—species. Functional Ecology 00, 1–12.

Sack L, Cowan P, Jaikumar N, Holbrook N. 2003. The ‘hydrology’of leaves: co-ordination of structure and function in temperate woody species. Plant, Cell & Environment 26, 1343–1356.

SIARL - Servizio Integrato Agrometeorologico della Regione Lazio. 2022. https://www.siarl-lazio.it/index.asp. Accessed May 2022.

Siefert A, Violle C, Chalmandrier L, et al. 2015. A global meta-analysis of the relative extent of intraspecific trait variation in plant communities. Ecology Letters 18, 1406–1419.

Silva MC, Teodoro GS, Bragion EFA, van den Berg E. 2019. The role of intraspecific trait variation in the occupation of sharp forest-savanna ecotones. Flora 253, 35–42.

Šmilauer P, Lepš J. 2014. Multivariate analysis of ecological data using CANOCO 5. Cambridge university press.

Solé-Medina A, Robledo-Arnuncio JJ, Ramírez-Valiente JA. 2022. Multi-trait genetic variation in resource-use strategies and phenotypic plasticity correlates with local climate across the range of a Mediterranean oak *(Quercus faginea)*. New Phytologist 234, 462–478.

Stefi AL, Christodoulakis NS. 2021. Approaching the “secrets” of the resin ducts in the mastic tree *(Pistacia lentiscus* L. cv. chia). Flora 285, 151940.

Thioulouse J, Dray S, Dufour A-B, Siberchicot A, Jombart T, Pavoine S. 2018. Multivariate analysis of ecological data with ade4. Springer.

UNEP. 1992. World atlas of desertification, vol. 80. UNEP and E. Arnold Ltd, Kent, UK.

Vignola C, Hättestrand M, Bonnier A, et al. 2022. Mid-late Holocene vegetation history of the Argive Plain (Peloponnese, Greece) as inferred from a pollen record from ancient Lake Lerna. PLOS ONE 17, e0271548.

Violle C, Navas M-L, Vile D, Kazakou E, Fortunel C, Hummel I, Garnier E. 2007. Let the concept of trait be functional! Oikos 116, 882–892.

Virta L, Teittinen A. 2022. Threshold effects of climate change on benthic diatom communities: Evaluating impacts of salinity and wind disturbance on functional traits and benthic biomass. Science of The Total Environment 826, 154130.

Welles SR, Funk JL. 2021. Patterns of intraspecific trait variation along an aridity gradient suggest both drought escape and drought tolerance strategies in an invasive herb. Annals of Botany 127, 461–471.

Wright IJ, Reich PB, Cornelissen JHC, et al. 2005. Modulation of leaf economic traits and trait relationships by climate. Global Ecology and Biogeography 14, 411–421.

Wright IJ, Reich PB, Westoby M, et al. 2004. The worldwide leaf economics spectrum. Nature 428, 821–827.

Zhang A, Zheng S, Didham RK, Holt RD, Yu M. 2021. Nonlinear thresholds in the effects of island area on functional diversity in woody plant communities. Journal of Ecology 109, 2177–2189.

Zhou J, Cieraad E, van Bodegom PM. 2022. Global analysis of trait-trait relationships within and between species. New Phytologist 233, 1643–1656.

